# Pericentriolar matrix integrity relies on cenexin and Polo-Like Kinase (PLK)1

**DOI:** 10.1101/2022.01.09.475500

**Authors:** Abrar Aljiboury, Amra Mujcic, Erin Curtis, Thomas Cammerino, Denise Magny, Yiling Lan, Michael Bates, Judy Freshour, Yasir H. Ahmed-Braimeh, Heidi Hehnly

**Affiliations:** Biology Department, Syracuse University, Syracuse NY

## Abstract

Polo-Like-Kinase (PLK) 1 activity is associated with maintaining the functional and physical properties of the centrosome’s pericentriolar matrix (PCM). In this study, we use a multimodal approach of human cells (HeLa) and zebrafish embryos in parallel to phylogenic analysis to test the role of a PLK1 binding protein, cenexin, in regulating the PCM. Our studies identify that cenexin is required for tempering microtubule nucleation and that a conserved C-terminal PLK1 binding site between humans and zebrafish is needed for PCM maintenance through mediating PLK1-dependent substrate phosphorylation events. PCM architecture in cenexin-depleted zebrafish embryos was rescued with wild-type human cenexin, but not with a C-terminal cenexin mutant (S796A) deficient in PLK1 binding. We propose a model where cenexin’s C-terminus acts in a conserved manner in eukaryotes, excluding nematodes and arthropods, to anchor PLK1 moderating its potential to phosphorylate PCM substrates required for PCM maintenance and function.

## RESULTS AND DISCUSSION

The centrioles and surrounding pericentriolar matrix (PCM) define the centrosome as one of the most complex non-membranous organelles in the cell [1,2]. Despite the centrosome’s structural and molecular complexity, the most characterized function of the centrosome is to nucleate and organize polarized microtubule arrays that generate cell polarity and form the structural framework for the mitotic spindle [1]. One way this function is regulated is by the centrosome acting as a scaffold to regulatory molecules, such as the mitotic kinase, Polo-Like Kinase 1 (PLK1). PLK1 is a major regulator of bi-polar spindle formation through PLK1-scaffold interactions at mitotic centrosomes/spindle poles. Once PLK1 is recruited to the centrosome, it modulates the phosphorylation and assembly of centrosome components, such as Pericentrin and Cep215, which are needed for γ-tubulin and γ-TuRC recruitment resulting in PCM expansion [3–5]. This expansion, termed centrosome maturation, is thought to play a crucial role in mitotic centrosome function during division [4,6]. Following maturation, continued PLK1 activity has been shown to be essential for PCM maintenance [7,8]. In zebrafish early embryos, pericentrin was identified to be delivered co-translationally to the centrosome [9] and then PCM maintenance at centrosomes likely requires PLK1 activity [10].

Two centrosome localized PLK1 scaffolds are Cep192 [11] and Odf2 isoform 9 called cenexin, [12–14]. Cep192 is reported to recruit an initial population of PLK1 and PCM components to centrosomes during bipolar spindle formation in mammalian, *Drosophila*, and *C. elegans* dividing cells [15–18]. However, much less is known about cenexin’s role during this time and the studies that have been performed were done primarily in murine or human cell models [12–14] with little known about cenexin’s conservation across phyla.

Previous studies identified a cenexin-dependent enrichment of PLK1 at the oldest mitotic centrosome in human cells [19]. Owing to the nature of centriole duplication, the two mitotic centrosomes that make up a spindle are inherently asymmetric from one another. The oldest (mother) mitotic centrosome is enriched with the centriole appendage protein cenexin, compared to the youngest mitotic centrosome (daughter) [20,21]. Cenexin is unique from other Odf2 isoforms in that it bears a C-terminal extension where the PLK1 binding site is situated [14]. Cenexin binds to PLK1 following Cyclin Dependent Kinase (CDK)1 phosphorylation of serine at position 796 (S796) [13]. However, the relationship between PLK1 and cenexin on the assembly and architecture of the PCM is unknown. Using a multidisciplinary approach, we found that cenexin’s C-terminus acts in a conserved manner in eukaryotes, excluding nematodes and arthropods, to anchor PLK1 moderating its potential to phosphorylate PCM substrates required for PCM maintenance and function.

### Cenexin loss results in PCM specific fragmentation

To examine the role of cenexin in PCM organization during metaphase, cells were depleted of cenexin using shRNA (cell lines reported in [22], Figure 1A). We noted significant loss of cenexin expression in cenexin shRNA treated cells (Figure 1A, S1A). We next examined changes in PCM organization in fixed cells where we immunostained for PCM components Cep192, pericentrin, Cep215, and γ-tubulin, along with a centriole marker centrin (Figure 1A-F). To identify centrosome organization defects, the 2-dimensional area of centrosome proteins was measured in cenexin depleted cells compared to control cells (Figure 1B-F). The centriole marker, centrin, and the PCM protein, Cep192, had no significant changes in centrosome area between cenexin shRNA and control cells (Figure 1A-1C), suggesting that the recruitment of these two proteins and/or maintenance at the centrosome is independent of cenexin. However, significant increases in Pericentrin, Cep215, and γ-tubulin area occurred in cenexin shRNA cells compared to controls (Figure 1A, 1D-F). This significant increase in PCM area was associated with significant increases in PCM fragmentation that were characterized as splayed or scattered (Figure S1B, example of scattered in Figure 1G). Strikingly, with scattered phenotypes we could find dramatic Cep215 puncta throughout the spindle, but centrin decorated centrioles still nicely positioned at the polar ends of the spindle (Figure 1G) and no significant defects in centriole number were noted (Figure S1C). Pericentrin and γ-tubulin fragments were consistently found to colocalize with one another (Figure S1D) and their fragmentation corresponded with defects in chromosome alignment (Figure 1G), a phenotype identified in [19]. These studies suggest that cenexin-loss is disrupting aspects of the PCM, involving Pericentrin, Cep215, and γ-tubulin, but not Cep192 or centriole number (Figure 1H).

**Figure 1.**
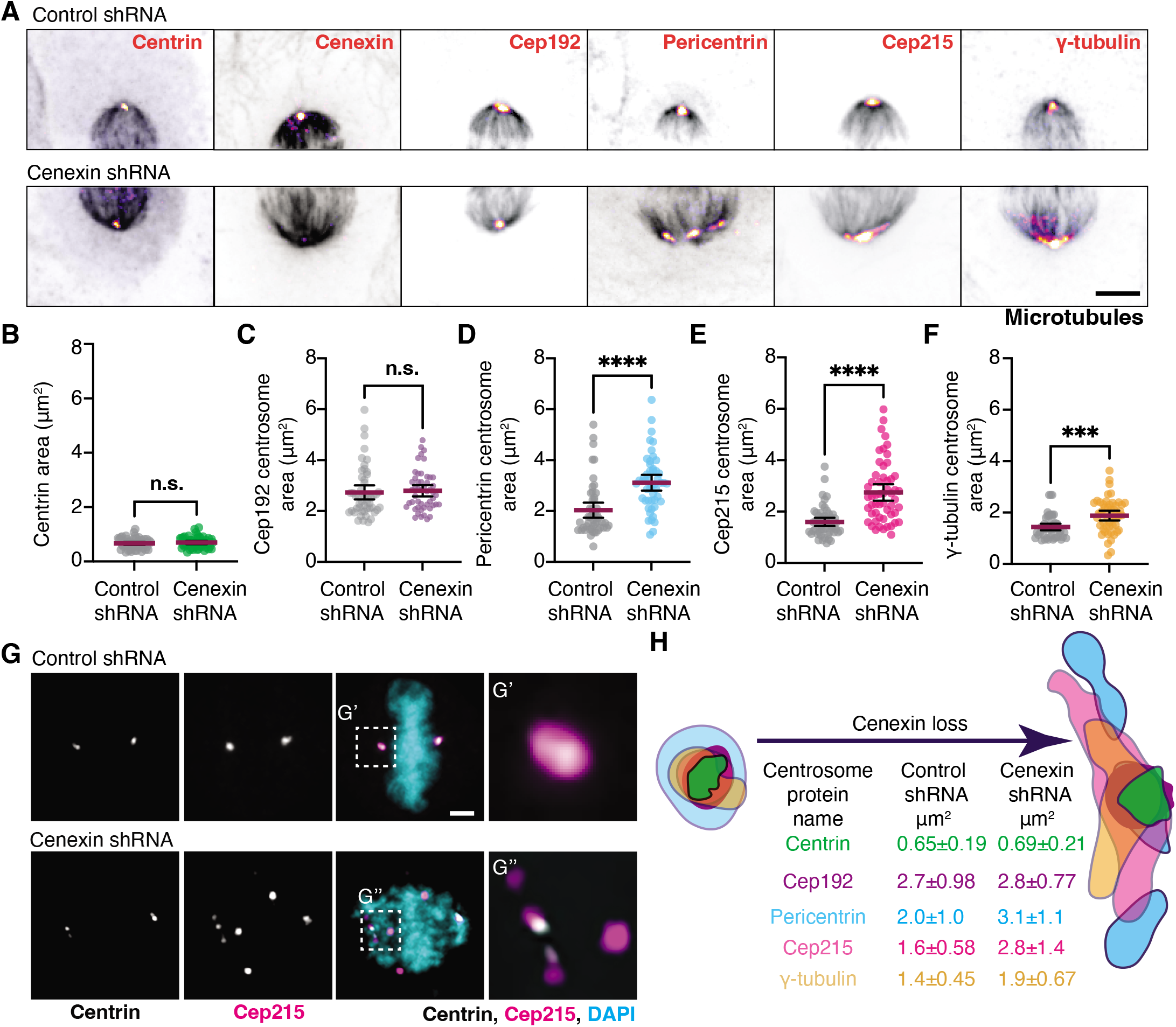
Cenexin loss results in PCM specific fragmentation. (A) Metaphase HeLa cells mitotic centrosomes labeled for centrosome markers: centrin, cenexin, Cep192, Pericentrin, Cep215 and γ-tubulin (Fire LUT) and microtubule marker, α-tubulin (grey). Control shRNA (top) and cenexin shRNA (bottom) treated cells shown. Scale bar, 5 μm. (B-F) Representative scatter plots depicting two-dimensional areas (μm^2^) of centrin (B), Cep192 (C), Pericentrin (D), Cep215 (E) and γ-tubulin (F) in control shRNA (grey) and cenexin shRNA. Mean (magenta) with 95% confidence intervals are displayed. Unpaired, two-tailed Student’s t-tests, n.s. not significant, ***p<0.001, ****p<0.0001. (G) Control shRNA (top) and cenexin shRNA (bottom) metaphase cell projection. Cells decorated with centrin (grey), Cep215 (magenta) and DNA (DAPI, cyan). Insets magnified 3x from G’ and G”. Scale bar, 5 μm. (H) Model depicting changes in PCM organization resulting from cenexin-loss. Colors corresponds with mean 2-dimensional areas (μm^2^) measured ±SD. For all graphs: detailed statistical analysis in Table S1. See also Figure S1.

To identify if there was any change in the amount of centrin, Cep192, Pericentrin, Cep215, and γ-tubulin recruited to the centrosome, a ratio of mean centrosome intensity of cenexin shRNA to control shRNA treated cells was calculated. If a ratio of 1 is obtained (grey dash line), it would indicate that there is no significant change in protein levels at the centrosome (Figure S1E). No significant deviation from 1 occurred for centrin, Cep192, Pericentrin, Cep215, and γ-tubulin (Figure S1E), suggesting the expanded area that Pericentrin, Cep215, and γ-tubulin occupy is unlikely due to increased protein abundance at the centrosome. We propose a model where the packing of Cep215, Pericentrin, and γ-tubulin complexes are compromised in cenexin depleted cells resulting in PCM fragmentation and an increase in overall area (Figure 1H).

### Cenexin tempers microtubule nucleation by mediating pericentrin associated acentrosomal nucleation sites

To determine whether cenexin loss and subsequent PCM disorganization (Figure 1) results in centrosome function defects, we performed a functional test to monitor mitotic centrosome–mediated microtubule nucleating activity over time (Figure 2A). Spindles were disassembled with nocodazole and examined at different times after nocodazole washout for microtubule nucleation (Figure 2A). At 0 min washout there was little to no detectable α-tubulin at mitotic centrosomes in both control and cenexin-depleted cells (Figure 2B). However, after 5 min of regrowth, mitotic centrosomes in cenexin-depleted cells demonstrated an increased ability to nucleate microtubules compared to control cells (Figure 2B-C). Intensity of α-tubulin was calculated at mitotic centrosomes, where a significant increase in signal was noted in cenexin depleted cells compared to control (Figure 2C). We also noted that an increased number of acentrosomal microtubule nucleation clusters occurred in cenexin depleted cells (8.0±0.502 sites) compared to control conditions 5 min post nocodazole washout (4.03±0.293 sites, Figure 2B, 2D). To examine when nucleation became elevated at the centrosome and when acentrosomal nucleation sites became significantly elevated post nocodazole washout we examined metaphase cells 30 sec, 1 min, 2 min, and 5 min post washout. Strikingly, nucleation was significantly elevated at mitotic centrosomes at each time point and was drastically more elevated at 5 min (4.34 times more than control, Figure S2A). This was consistent with acentrosomal nucleation sites marked by α-tubulin clusters, where at 5 min twice as many clusters were observed in cenexin-depleted cells to control cells (Figure S2B). While control and cenexin-depleted cells had acentrosomal microtubule clusters at 5 min, with cenexin-depleted cells containing more, we found that by 20 min a nicely formed bipolar spindle still occurred (Figure 2B). This suggests that the acentrosomal microtubule clusters that form can transport to the main mitotic centrosomes in both control and cenexin-depleted cells to assist in the formation of a bipolar mitotic spindle [23,24], but the presence of more acentrosomal microtubule clusters in cenexin-depleted cells is likely causing the significant elevation in microtubule nucleation.

**Figure 2.**
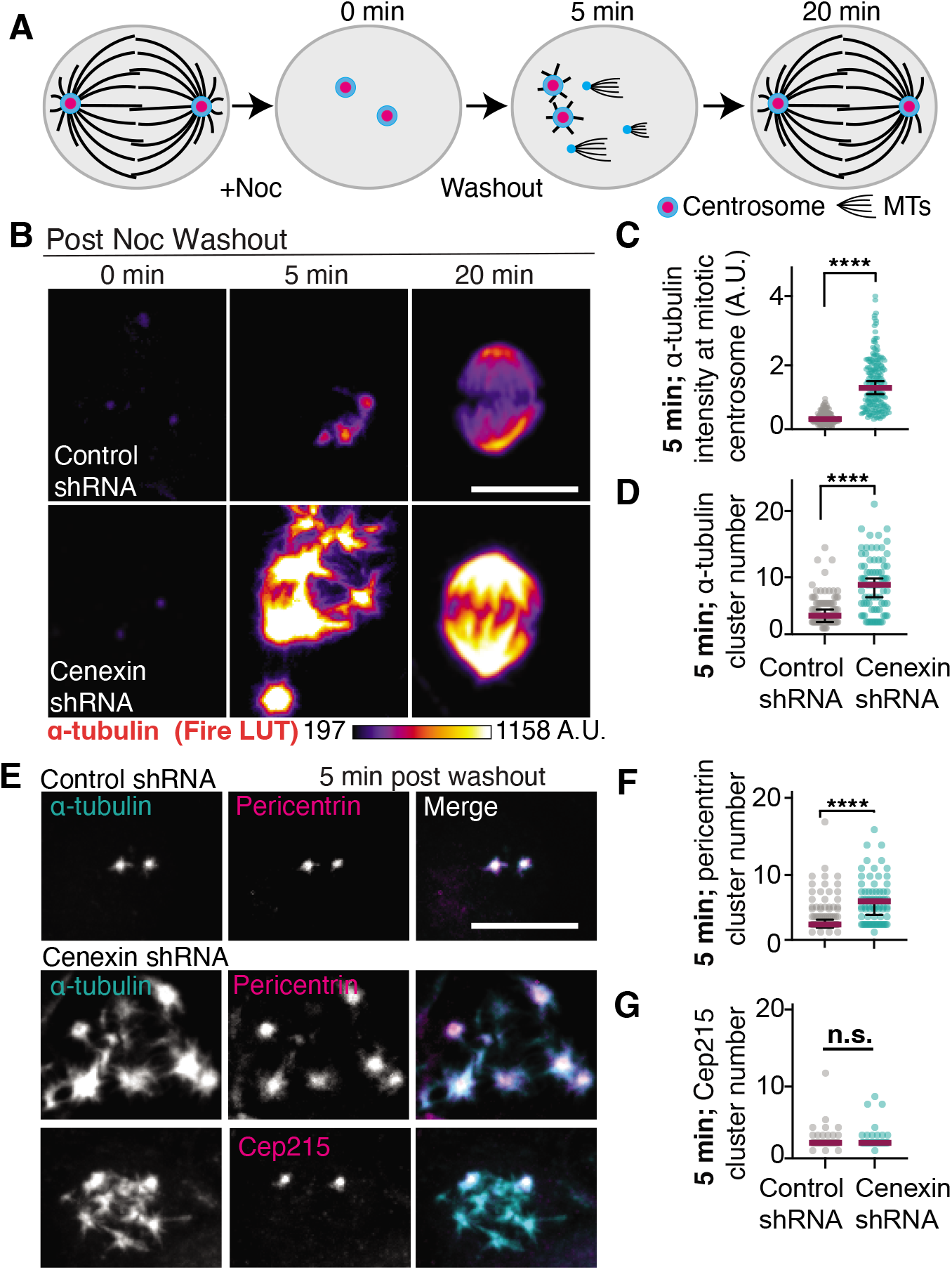
Cenexin tempers microtubule nucleation by mediating pericentrin associated acentrosomal nucleation sites. (A) Model of microtubule renucleation assay in metaphase cells. (B) Control and cenexin shRNA treated HeLa cells are incubated with nocodazole then washed with fresh media (washout). At 0 min, 5 min and 20 min post washout, cells are fixed and immunostained for α-tubulin (Fire LUT). Scale bar, 10 μm. (C-D) Scatter plot depicting α-tubulin intensity at metaphase centrosomes (C) or the number of α-tubulin clusters (D) 5-min post nocodazole washout in control shRNA (grey) and cenexin shRNA (cyan). Mean (magenta) with 95% confidence intervals are displayed. Unpaired, two-tailed Student’s t-tests, ****p<0.0001. (E) α-tubulin (grey, cyan in merge), pericentrin (gray; magenta in merge) and CEP215 (gray; magenta in merge) metaphase cell projections in control shRNA or cenexin shRNA at 5-min post nocodazole washout. Scale bar, 10 μm. (F-G) Scatter plot depicting the number of pericentrin clusters (F) and CEP215 clusters (G) 5-min post nocodazole washout in control shRNA and cenexin shRNA cells. Mean (magenta) with 95% confidence intervals are displayed. Unpaired, two-tailed Student’s t-tests, ****p<0.0001, n.s. not significant. For all graphs: detailed statistical analysis in Table S1. See also Figure S2.

We next examined whether two PCM components that are known to form a complex through direct interaction, Cep215 and pericentrin [6,25], localized with α-tubulin at acentrosomal microtubule clusters in cenexin-depleted cells compared to control. To test for localization of Cep215 and pericentrin with α-tubulin we calculated a Mander’s overlap coefficient to measure the degree of colocalization (0 to 1, with 1 being perfectly colocalized) from metaphase cells 5 min post nocodazole washout. In control cells, we found that Cep215 and pericentrin colocalized with α-tubulin 5 min post nocodazole washout (above 0.5 coefficient, Figure S2C-D)). However, with cenexin-loss pericentrin maintained its localization with α-tubulin, whereas Cep215 did not colocalize to the same extent (0.41 coefficient, Figure S2C-D). Along these same lines, we find that pericentrin associates with α-tubulin clusters/acentrosomal sites in cenexin-depleted cells 5 min post washout, whereas Cep215 is not present at these sites (Figure 2E). Consistent with this, cenexin-depleted cells on average have 5.38±0.354 pericentrin clusters compared to 2.30±0.122 Cep215 clusters (Figure 2F-G). This finding suggested that Cep215 and pericentrin can no longer maintain a stable association with cenexin loss and pericentrin is associated more so to the acentrosomal microtubule clusters.

### Cenexin and PLK1 work together to maintain proper PCM organization

We examined whether PCM organization is modulated by cenexin through its identified role as a PLK1 binding partner [12,13]. Specifically, we wanted to examine if cenexin loss causes changes in active-PLK1 at centrosomes and/or PCM substrate phosphorylation that associates with PCM fragmentation. To test for PCM substrate phosphorylation we immunostained for phosphorylated serine/threonine (pS/T) modified proteins in relation to the PCM using confocal (Figure 3A) or expansion microscopy (ExM, Figure 3D). ExM is an imaging protocol which provides sub-diffraction limited details of centrosome organized components [26–28]. Using quantitative confocal microscopy, cenexin-depleted mitotic cells demonstrated elevated PCM substrate phosphorylation (pS/T) compared to control conditions (Figure 3A, 3B). The elevated phosphorylation state found in cenexin-depleted cells was alleviated when cells were treated with a PLK1 small molecule inhibitor, BI2536 (Figure 3A, 3B), suggesting that elevated pS/T was specific to PLK1. Using ExM, we found that pS/T and Pericentrin modified centrosome substrates encompassed a significantly larger area in cenexin depleted cells (16.68±4.99 μm^2^ for pS/T, and 41.46±13.64μm^2^ for Pericentrin) compared to control cells (9.72±3.35 μm^2^ for pS/T, and 12.92±4.17 μm^2^ for Pericentrin, Figure 3D-F). Notably, we identified that cenexin-depleted cells treated with BI2536, had normal PCM area measured by either Cep215 in non-ExM cells (Figure S3A-B) or pericentrin in ExM cells (Figure 3D, 3F). ExM also provided insight that we likely wouldn’t obtain from traditional confocal techniques. By expanding the PCM from approximately 1.47±0.07 μm^2^ under control conditions (Figure S3B) to 12.92±0.81 μm^2^ with ExM (Figure 3F), we were able to identify that pericentrin always occupies a significantly larger area than pS/T both in control (12.92±4.17 μm^2^ to 9.72±3.35 μm^2^) and cenexin-depleted cells (41.46±13.64μm^2^ to 16.68±4.99 μm^2^), suggesting pS/T substrates have a concentrated distribution towards the centrosome center in relation to pericentrin (Figure 3D-F, 3G). This tiered organization is not necessarily perturbed by cenexin-depletion, but cenexin depletion expands PCM size and pS/T distribution and frequency (Figure 3G). These studies suggest that cenexin-depleted mitotic cells have fragmented PCM potentially due to elevated PLK1-dependent phosphorylation events on PCM substrates.

**Figure 3.**
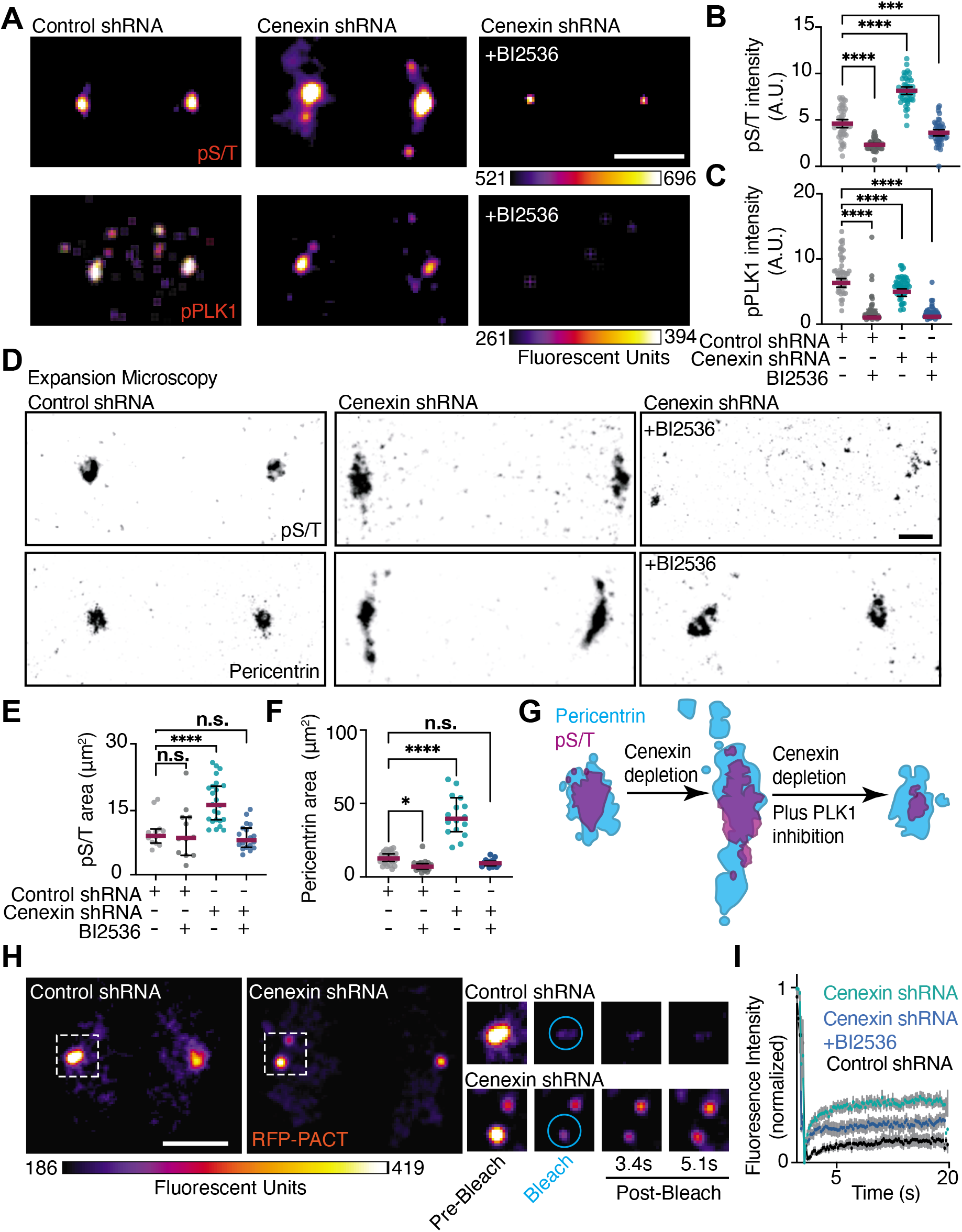
Cenexin and PLK1 work together to maintain proper PCM organization. (A) Control shRNA or cenexin shRNAs metaphase cells with or without BI2536 were immunolabeled for phospho-serine/threonine (pS/T, top) and phospho-PLK1(T210) (pPLK1, bottom) (FireLUT). Scale bar, 5μm. (B-C) Representative scatter plots depicting pS/T (B) or pPLK1 intensity (D) at metaphase centrosomes. Mean (magenta) with 95% confidence intervals are displayed. One-way ANOVA with multiple comparisons to control cells, ****p<0.0001. (D) Expansion microscopy of control and cenexin shRNA metaphase cell centrosomes with or without BI2536. Centrosomes immunolabeled for pS/T (inverted grey, top) or Pericentrin (inverted grey, bottom). Scale bar, 5μm. (E-F) Representative scatter plots depicting expanded two-dimensional areas (μm^2^) of pS/T (E) or Pericentrin (F) in control shRNA and cenexin shRNA metaphase cells with or without BI2536. Mean (magenta) with 95% confidence intervals are displayed. One-way ANOVA with multiple comparisons to control cells, n.s. not significant, **p<.01, ****p<0.0001. (G) Model depicting relative changes in pS/T and Pericentrin organization with cenexin depletion and cenexin depletion coupled with PLK1 inhibition. (H) FRAP examples of control and cenexin shRNA metaphase cell centrosomes. Magnified insets of boxed centrosomes depict examples of pre-bleach, bleach, 3.4s post-bleach and 5.1s post-bleach conditions in control and cenexin shRNA cells. Cyan circles represent region to which 405nm laser was applied for photo-bleaching. Scale bar, 5 μm. (I) FRAP curves displaying changes in normalized RFP-PACT signal overtime (s) in metaphase control (black), cenexin shRNA (cyan) and cenexin shRNA with BI2536 cells (blue). SEM shown. For all graphs: detailed statistical analysis in Table S1. See also Figure S3.

We tested if the elevated amounts of pS/T in cenexin-depleted cells resulted from either too much PLK1 at the centrosome or an overabundance of active PLK1 at the centrosome by immunostaining for PLK1 (Figure S3A) and active phospho-PLK1(T210) (Figure 3A) in the presence or absence of the PLK1 small molecule inhibitor (BI2536). BI2536 treatment resulted in a robust and significant decrease in pPLK1(T210) (Figure 3A, 3C) in both control and cenexin-depleted cells, demonstrating that BI2536 was specifically decreasing PLK1 activity. We identified a slight, but significant decrease in PLK1-activity (pPLK1(T210) at mitotic centrosomes (Figure 3A, C) compared to total PLK1 (Figure S3A, C) in cenexin-depleted cells compared to control (Figure 3C). We were surprised to observe nearly similar levels of active pPLK1(T210) between control and cenexin-depleted conditions since we saw elevated substrate phosphorylation levels in cenexin-depleted cells (Figure 3B) and expected to find elevated active pPLK1(T210) levels. One potential interpretation from our findings is that under cenexin-depleted conditions a small population of normally cenexin-bound pPLK1(T210) is now free to act on PCM substrates resulting in PCM fragmentation in cenexin-depleted cells.

We addressed if the biophysical state of the PCM was altered with loss of cenexin. A PCM marker, RFP-pericentrin AKAP450 C-terminal centrosome targeting domain (PACT), was expressed in control and cenexin-depleted cells treated with and without the PLK1 inhibitor BI2536 (Figure 3H-I). Using fluorescence recovery after photobleaching (FRAP) we examined the mobility of RFP-PACT at metaphase centrosomes, where we found significantly different dynamics between cenexin-depleted cells and control (Figure 3H-I, S3D). Control cells mean mobile fraction (0.15±0.04) was significantly less than the mobile fraction of cenexin depleted cells (0.35±0.15, Figure 3H-I, S3D). This mobility was partially rescued in cenexin-depleted cells treated with the PLK1 inhibitor BI2536 (0.23±0.12, Figure 3H-I, S3D). Together these studies suggest that the increased substrate phosphorylation through PLK1-cenexin disruption is likely causing increased PCM mobility. This increase in PCM mobility is suggestive of a model where the packing of Pericentrin is compromised in cenexin-depleted cells that ultimately results in its fragmentation (Figure 3G).

### Cenexin phosphorylation at its conserved C-terminal PLK1 binding site is required for maintenance of PCM in vivo

Human cenexin (Odf2 isoform 9) is unique from other human Odf2 isoforms in that it bears an N- and C-terminal extension (1-42 amino acid stretch for N-terminus, 701-805 amino acid stretch for C-terminus), with the PLK1 binding site being situated within the C-terminus [14] (Figure 4A, 4C). Previous studies identified that human cenexin binds to PLK1 following CDK1 phosphorylation of S796 [13]. Here, we examined whether human cenexin, and its N- and C-terminal extensions, are present across phyla by performing BLASTp searches of the human sequences against 67 proteomes through NCBI’s webserver (Figure 4B). Since sequence hits for several species were not detected on the web server due to insufficient similarity, we performed BLASTp searches using a local BLAST installation to identify possible orthologous sequences (Figure S4A). With these studies, we found that cenexin is present in all vertebrates (chordates) along with two other centrosome proteins tested, the PLK1 scaffold CEP192 [18], and the centriole protein centrin [29,30]. Cenexin is also present in hemichordates, mollusks, annelids, rotifers, platyhelminths, and cnidarias, but absent in arthropods and nematodes, along with ctenophores, poriferas, and placozoa. The presence of cenexin in each phylum requires the presence of centrioles (ex. chordates), however the presence of centrioles does not necessarily guarantee that cenexin is present (ex. nematodes, Figure 4B, S4C). When performing BLASTp searches through the webserver, the C-terminal part of cenexin is conserved in all vertebrates (chordates, excluding genus pongo, Figure 4B), hemichordates, molluscs, annelids, rotifers, platyhelminths, and cnidarias, whereas the N-terminus is only present in chordates (excluding genus pongo, danios, and petromyzon) and hemichordates (Figure 4B). However, using a local BLAST installation pongo and danio was identified with high-confidence to include both the N- and C-terminus, but oryctolagus along with petromyzon excludes the N-terminus (Figure S4C). These findings suggest that the C-terminus containing the PLK1 binding site evolved first and is highly conserved in its function and potential regulation of the centrosome across phyla. Interestingly, our analyses reveal that the PLK1 scaffold Cep192 [18,31], is detected in arthropods where the cenexin C-terminal extension is lacking. Additionally, a Cep192 “like” centrosome protein, spd-2, that acts as a PLK1 scaffold was identified in *C. elegans* that was not identified by BLASTp analysis [16]. This suggests that Cep192 may have some redundancy in PLK1 scaffolding functions allowing cenexin to be potentially dispensable in arthropods and *C. elegans* (Figure 4B).

**Figure 4.**
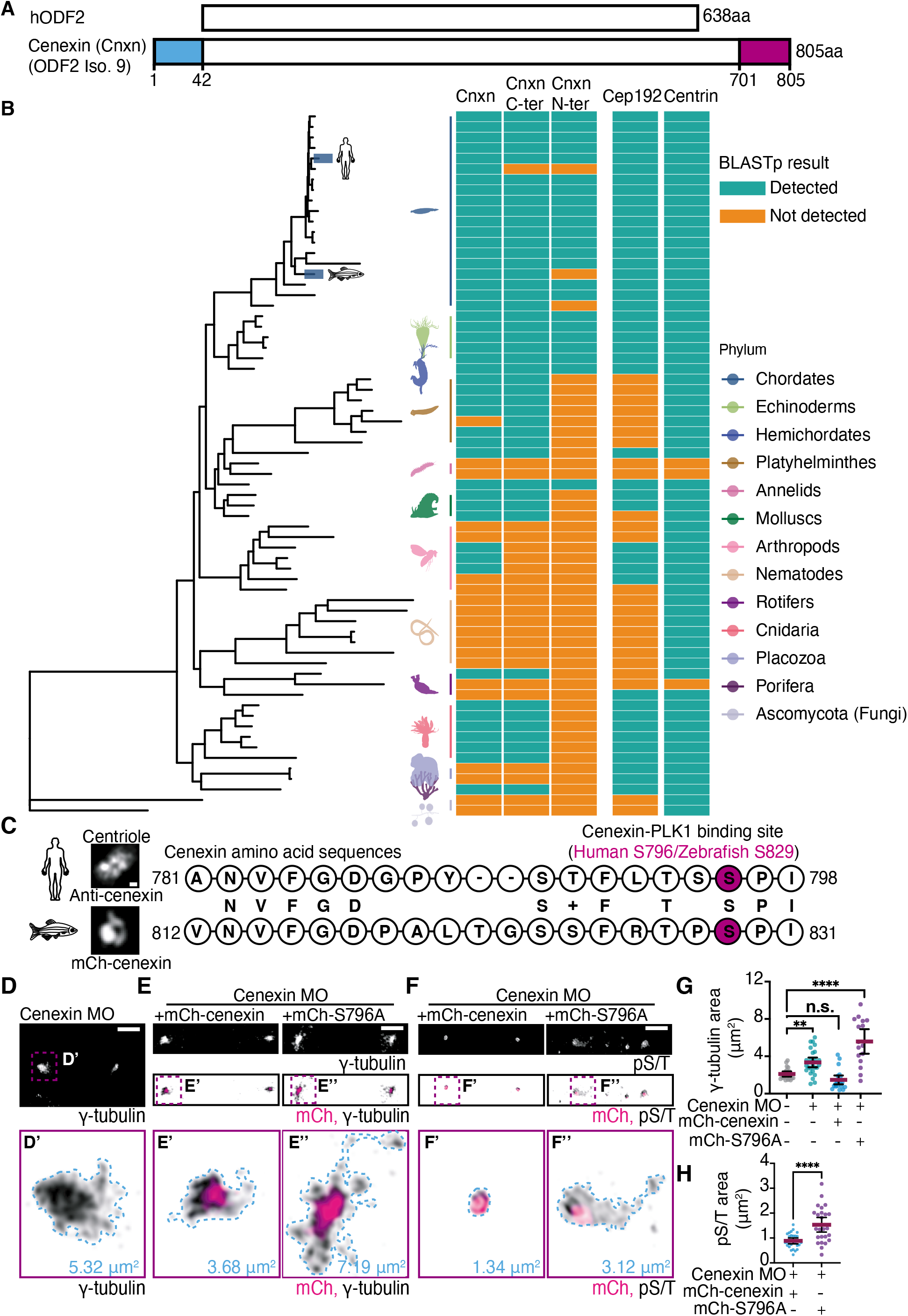
Cenexin phosphorylation at its conserved C-terminal PLK1 binding site is required for maintenance of PCM *in vivo*. (A) Schematic representation of human hODF2 and cenexin (Cnxn, ODF2 isoform 9). The blue and magenta boxes highlight the N- and C-terminal extensions unique to cenexin. (B) Phylogenetic tree of the evolution of cenexin and its N-terminus and C-terminus in relation to Cep192 and centrin across different animal phyla. Cyan and orange boxes indicate whether cenexin and centrosome components (Cep192 and centrin) are detected or not detected within representative species of each phylum. Dark blue boxes highlight two species (humans and zebrafish). (C) Amino acid alignment between human and zebrafish cenexin C-terminal PLK1 binding motif. The serine highlighted in magenta represents the known human cenexin-PLK1 site (S796) and potential zebrafish biding site (S829). Letters in the middle between two sequences represent identical amino acids, and the + sign represent an amino acid of functional identity. Representative confocal maximum projection of cenexin from expanded (ExM) human cells and from a zebrafish embryo cell shown. Scale bar 0.05 μm. (D-F) Representative cells from 512-cell zebrafish embryos under cenexin depletion conditions (Cenexin MO, D), or rescue conditions (cenexin MO plus mCh-cenexin or mCh-cenexin-S796A, magenta in E-F) fixed and immunostained for γ-tubulin (inverted grey, D-E) or pST (inverted grey, F). Insets (D’, E’, E”, F’, and F”) at 5x magnification, corresponding areas outlined (μm^2^). Scale bar, 5μm. (G) Scatter plot depicting γ-tubulin area (μm^2^) at mitotic centrosomes under control, cenexin MO, and rescue conditions (cenexin MO plus mCh-cenexin or mCh-cenexin-S796A). Mean (magenta) with 95% confidence intervals shown. One-way ANOVA with multiple comparisons to control cells, n.s. not significant, **p<.01, ****p<0.0001. (H) Scatter plot depicting pS/T area (μm^2^) at mitotic centrosomes under rescue conditions (cenexin MO plus mCh-cenexin or mCh-cenexin-S796A). Unpaired, two-tailed Student’s t-tests, ****p<0.0001. For all graphs: detailed statistical analysis in Table S1. See also Figure S4.

To determine whether the role cenexin plays in PCM organization is conserved between human cells and an *in vivo* chordate model identified to have the C-terminal extension, we aligned the amino acid sequence of human cenexin to zebrafish (*Danio rerio*) cenexin (Figure 4B, 4C). The two sequences shared 50.36% identity with a 70% positivity rate that accounts for amino acids with functional similarity, suggesting high conservation between human and zebrafish cenexin. Human cenexin amino acids spanning the S796 site (781-798) were aligned to zebrafish cenexin where S829 corresponded with human S796 (Figure 4C). This provides further support that the C-terminal tail containing the PLK1 binding site has a conserved function across species. To test this, we depleted cenexin from zebrafish embryos using previously published translational morpholinos (MO) to block the translation of cenexin mRNAs [32] and rescued with human wild-type cenexin (mCherry-cenexin) or cenexin deficient in PLK1 binding (mCherry-cenexin-S796A). mCherry-cenexin displayed a compact ring-like centriole organization at zebrafish 512-cell stage centrioles comparable to immunostaining for endogenous cenexin using ExM in human cells (Figure 4C, S4B). Both mCherry-cenexin and -cenexin-S796A had concentrated organization towards the centrosome center (Figure 4E) with no significant difference in centrosome area occurring between the two (Figure S4C). When depleting endogenous zebrafish cenexin, γ-tubulin became more fragmented and occupied a significantly larger area (5.32 μm^2^ in Figure 4D, 4G) compared to control conditions (Figure 4G) or wild-type cenexin (mCherry-cenexin) rescue conditions (3.68 μm^2^ in Figure 4E, 4G). However, mCherry-cenexin-S796A was unable to rescue centrosome area (7.19 μm^2^ in Figure 4E, 4G), suggesting that the ability of cenexin to bind to PLK1 at S796 is required for its function to modulate PCM organization.

These results prompted us to examine whether PCM substrate phosphorylation relies on cenexin *in vivo*. To do this, we fixed and immunostained cenexin depleted embryos rescued with mCherry-cenexin or mCherry-cenexin-S796A for pS/T modified proteins in relation to the PCM (Figure 4F, 4H). With mCherry-cenexin rescue conditions pS/T signal was tightly organized (0.88±0.30 μm^2^, Figure 4F, 4H) occupying a smaller centrosome area than mCherry-cenexin (1.36±0.94 μm^2^, Figure S4A). Whereas with mCherry-cenexin-S796A rescues, pS/T signal was mis-organized and dispersed (1.5±0.71 μm^2^ compared to controls at 0.88±0.30 μm^2^, Figure 4F, 4H). These results are in agreement with what we found *in vitro* (Figure 3G), where pS/T substrates have a concentrated distribution towards the centrosome center in relation to the PCM that is noted by either Pericentrin in human cells (Figure 3D-F, 3G) or γ-tubulin in zebrafish (Figure 4E-F). This tiered organization is not perturbed by loss of cenexin-PLK1 binding, but inhibition of this interaction results in expanded pS/T distribution (Figure 4F, 4H). Altogether, our experiments indicate that the conserved PLK1 binding motif in the C-terminal tail of cenexin is required for PCM maintenance and function.

## Supporting information

Supplemental figures and stats

## ACKNOWLEDGEMENTS

This work was supported by National Institutes of Health grants R01GM127621 (H.H.) and R01GM130874 (H.H.). This work was supported by the U.S Army Medical Research Acquisition Activity through the FY16 Prostate Cancer Research Programs under Award no. W81XWH-20-1-0585 (H.H.). Opinions, interpretations, conclusions, and recommendations are those of the authors and not necessarily endorsed by the Department of Defense.

## AUTHOR CONTRIBUTIONS

H.H., A.A.A., A.M., and E.C. designed, performed, and analyzed experiments; H.H. and A.A.A. wrote manuscript; Y.A.-B. built phylogenetic tree; J.F. and M.B. provided molecular reagents and zebrafish husbandry. D.M., T.C., and Y.L. performed experiments and associated analyses. All authors provided edits. H.H. oversaw project.

## METHODS

### RESOURCE AVAILABILITY

#### Lead Contact

For further information or to request resources/reagents, contact the Lead Contact, Heidi Hehnly.

#### Materials Availability

No new materials were generated for this study.

#### Data and Code Availability

All data sets analyzed for this study are displayed. A supplemental CSV file is provided with the species used in the phylogenetic analysis with a FTP link to download their proteomes (supplemental file SF1A, Metazoan_taxon_list_FTP_links.csv), the tree file is provided in Newick format (File SF1B, nuclear_genes_tree.nwk), the R script file for phylogeny analysis is available as a supplemental file SF1 (File SF1C, phylo_analysis.html), and the BLAST results are provided in a supplement table formatted in supplemental file SF1 (File SF1D, NCBI_BLAST_results.zip; File SF1E, Homebrew_BLAST_results.zip).

### EXPERIMENTAL MODEL AND SUBJECT DETAILS

#### Zebrafish

All zebrafish lines were maintained using standard procedures approved by the Syracuse University IACUC committee (protocol #18-006). For details see [10,33]. See Key Resources Table for list of zebrafish transgenic lines used.

#### Cell Culture

HeLa cells treated with either control or cenexin shRNA [22] were used throughout this study. Cells were selected in puromycin (3μg/mL). See Key Resources Table for list of cell lines used.

### METHOD DETAILS

#### Immunofluorescence

Cells were plated on #1.5 coverslips until they reached 90% confluence and fixed using ice cold methanol for 10 min. Standard immunostaining procedures were performed (described in [34]). Coverslips were rinsed with diH2O and mounted on glass slides using ProLong Gold mounting media (Thermo Fisher Scientific; P36934). Zebrafish embryos were fixed using 4% PFA at 4°C overnight, for immunostaining see [10,33]. See Key Resources Table for list of antibodies used in this study.

#### Chemical Inhibitors

Chemical inhibitors include nocodazole used on cells at 100 nM or 10μm, ProTAME (Fisher; I44001M) used on cells at 10 μM, and BI2536 used on cells at 100 nM. See Key Resources Table for more information on inhibitors used in this study.

#### Morpholino Injections

Anti-cenexin vivo translational morpholinos (Gene Tools; [32]) or vivo standard control morpholinos (Gene Tools) were constituted as 1mM stock in water and injected into zebrafish yolks at 1 cell stage in a final concentration of 2ng/nL. Injection protocols detailed in [33].

#### Plasmid Constructs and mRNA

Gibson cloning methods were used to generate mCherry-cenexin-WT and mCherry-cenexin-S796A plasmids (NEBuilder HiFi DNA assembly kit), then purified using DNA maxi-prep kit (Bio Basic; 9K-006-0023). mRNA was generated from plasmids using mMESSAGE mMACHINE^™^SP6 transcription kit (Thermo Fisher Scientific; AM1340).

#### Fluorescence Recovery After Photobleaching (FRAP)

FRAP experiments were performed using a Leica DMi8 STP800 spinning disk confocal microscope using a 40x/1.10 NA water immersion objective. A Region of Interest (ROI) was placed over one of the spindle poles, where a 405nm laser was applied. RFP-PACT fluorescence was bleached within the ROI after administration of the laser. Fluorescence recovery time as well as signal intensity were measured every 100 ms to determine the mobile fraction.

#### Microtubule Renucleation Assay

HeLa cells were treated with 1 μM nocodazole in media for 30 minutes. Cells were washed twenty times with 1xPBS and placed in media at 37 °C for 0s, 30s, 1m, 2m, 5m or 20m. Cells were fixed using methanol overnight at −20 °C. Cells were immunostained and then imaged for analysis.

#### Expansion Microscopy

Protocol used was modified from previously published expansion protocols [26,35]. Modified protocol is described below. See Key Resources Table for chemicals used.

##### Cell preparation

HeLa cells were grown on glass coverslips and synchronized with 10 μM protame for 1 hour, then fixed with ice cold methanol for 10 minutes or 4% PFA in 1 X PBS for 1 hour at 22-25 °C.

##### Immunostaining and acrylamide incubation

Cells were blocked in PBSΔT for 1 hour at 22-25 °C if methanol fixed or in 0.1% Triton X-100 with 5% donkey serum in 1x PBS for 15 minutes at 22-25 °C if PFA fixed. Cells were then incubated with primary antibodies in blocking buffer overnight at 4 °C then with secondary antibodies for 4 hours at 22-25 °C. Cells were incubated in 30% acrylamide solution in 1xPBS overnight at 40 °C.

##### Gelation, cell punching, and digestion

Following three 10-minute 1 X PBS washes (PFA fixed) or 10x PBSΔT washes (if methanol fixed), the slides (cell side up) and the parafilm-covered Petri dish were placed on an ice bath. Gelation reagents were placed on the coverslips in chilled 1 X PBS: 20% acrylamide, 7% sodium acrylate, and 0.04% bisacrylamide, with APS and TEMED added just before application. Gelation solution was added to each coverslip, incubated for 20 minutes on ice, and then incubated at 30°C for 1.5 hours. A 4 mm biopsy punch was utilized to excise punches from gelled samples. Punches were incubated with digestion buffer (Triton-X + EDTA + Tris pH8 + NaCl in water) overnight at 22-25 °C in the dark.

##### Post expansion staining to enhance fluorescence

Cell punches were washed and blocked with appropriate buffer then incubated with primary antibodies followed by secondary antibodies in blocking buffer for 4 hours at 22-25 °C.

##### Expansion and mounting

After digestion, samples were expanded in dH2O for 2 hours at 22-25 °C with dH2O exchange every 20 minutes. Samples were left in dH2O and DABCO overnight to expand and protect IF signal. Samples were mounted in MatTek plate the following day for imaging.

#### Imaging

Tissue culture cells and zebrafish embryos were imaged using a Leica DMi8 STP800 (Leica, Bannockburn, IL) equipped with X-light V2 Confocal Unit spinning disk and an 89 North-LDI laser launch with a Photometrics Prime-95B camera or a Leica SP8 Laser Scanning Confocal Microscope (LSCM; Leica, Bannockburn, IL). Optics used on Leica DMi8 were HC PL APO 63x/1.40 NA oil CS2 or HC PL APO 40x/1.10 NA CORR WCS2 water. The optics used on the SP8 LSCM is HC PL APO 40x/1.10NA CORR CS2 0.65 water objective or HC PL APO 63x/1.30 NA Glyc CORR CS2 glycerol objective. Visiview software (Lecia DMi8) or LasX (SP8 LSCM) were used to acquire images.

For live Zebrafish imaging, fluorescent transgenic or mRNA injected embryos (injection protocols in [33]) were mounted in 2% low melting agarose gel (Thermo Fisher Scientific; 16520100) at 512-cell stage and imaged using SP8 LSCM or spinning disk confocal microscopes.

#### Image and Statistical Analysis

Images were processed using FIJI/ImageJ software. All analysis was performed on maximum projections unless otherwise stated. Spindle pole area was measured using the freehand selection tool to draw a boundary of the poles and calculating the area within shape [33]. Signal intensities were measured by placing a ROI over site of interest (poles or clusters). Signal intensity was calculated by subtracting the minimum intensity from the mean intensity of measured region, unless otherwise noted. All figures were created in Adobe Illustrator, and graphs were created using Graphpad Prism software. Statistical analyses (unpaired Student’s t tests and analysis of variance ANOVA) were performed using Graphpad Prism. **P<0.01, **P<0.001, and ***P<0.0001.

#### Phylogenetic Analysis of Cenexin

To analyze the pattern of evolutionary conservation of Cenexin (Cnxn, Odf2 isoform 9) among metazoan phyla, a metazoan phylogeny was built using a single protein isoform (isoform 1) of three nuclear genes (*SMCA1A*, *SMC1B*, *and MCM5*) for 67 species that represent the major metazoan phyla and for which annotated proteomes are available on NCBI (supplemental files SF1A-B). *Saccharomyces cerevisiae* and *Neurospora crassa* were used as outgroups. The three protein sequences were first extracted from the *Homo sapiens* proteome and orthologs in the remaining species were identified using BLASTp [36]. A multiple sequence alignment of the three proteins was constructed using Muscle [37], and individual alignments were subsequently concatenated. A maximum likelihood phylogenetic tree was built with PhyML using default parameters [38]. To identify orthologs of Cenexin and its N- and C- termini, we first performed BLASTp searches of the human sequences against the 67 proteomes on the NCBI web server. Sequence hits for several species were not detected on the web server due to insufficient similarity, therefore we performed BLASTp searches using a local BLAST installation to identify possible orthologous sequences. Because we obtained a BLASTp hit for each species, we classified hit confidence as potential orthologs based on hit length, percent identity, and NCBI annotation. Specifically, we considered a hit a “high confidence” ortholog if the protein was annotated by NCBI as an Odf2 ortholog or had a matched alignment length >100 a.a. (Cnxn), >80 a.a. (C-terminus), or >30 a.a. (N-terminus) and a percent identity >60% (Cnxn) or >40% (C-terminus and N-terminus). Alternatively, we considered a hit a “medium confidence” ortholog if the matched alignment length was between 30 a.a. and 100 a.a. (Cnxn), 30 a.a. and 80 a.a. (C-terminus), or 40 a.a. and 30 a.a. and a percent identity >40%. Finally, alignment matches that were shorter than 30 a.a. (Cnxn and C-terminus) or 15 a.a. (N-terminus) and percent identity <40% were considered “low confidence” hits. Any hit that did not match our criteria was considered an “unlikely ortholog”. All phylogenetic analyses were performed on R primarily using the ggtree package [39]. The phylogenetic analysis script and input files is available in supplemental files SF1C-E.

## KEY RESOURCES TABLE

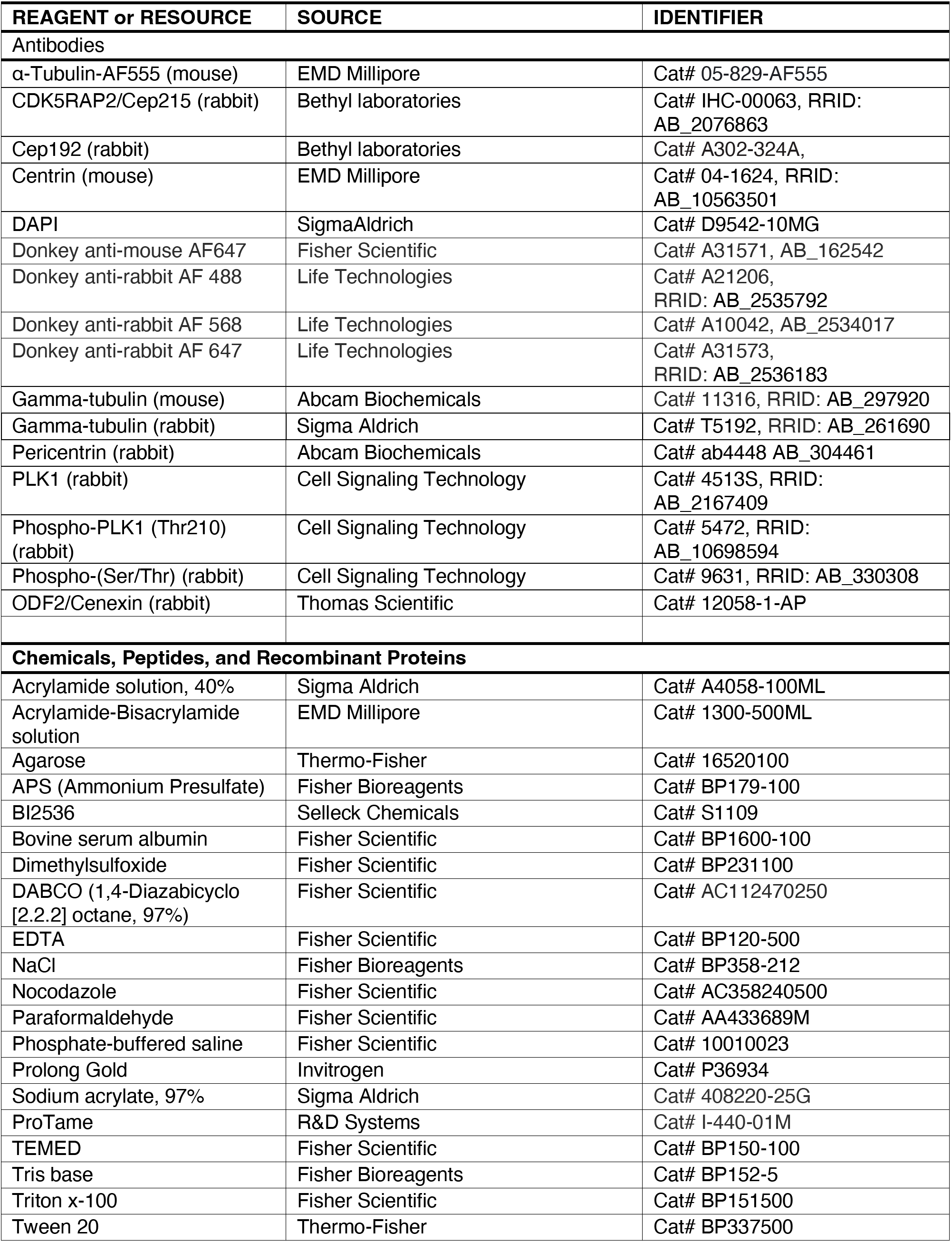
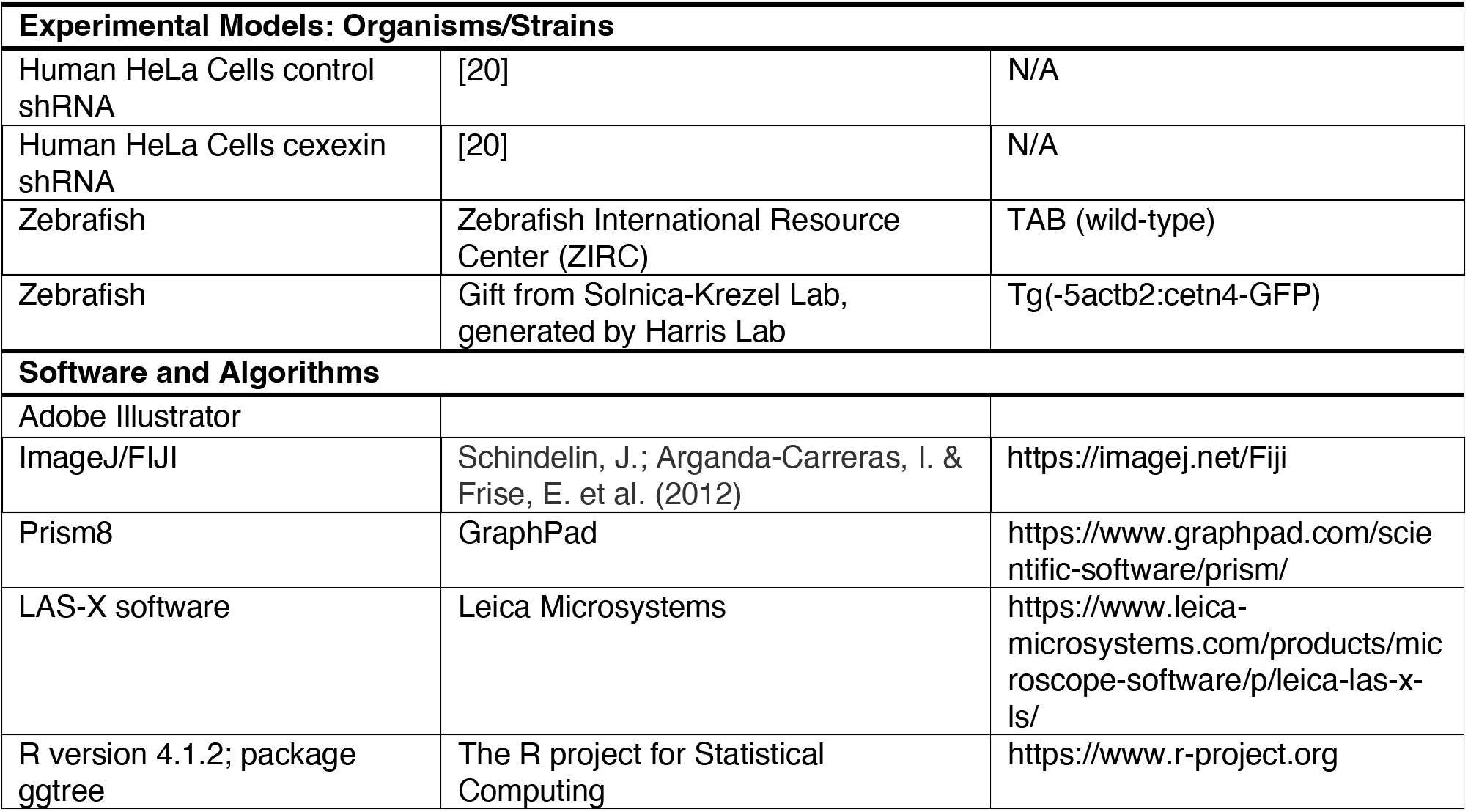

## REFERENCES

1. Vertii, A., Hehnly, H., and Doxsey, S. (2016). The centrosome, a multitalented renaissance organelle. Cold Spring Harb. Perspect. Biol. 8.

2. Mennella, V., Agard, D.A., Huang, B., and Pelletier, L. (2014). Amorphous no more: subdiffraction view of the pericentriolar material architecture. Trends Cell Biol. 24, 188–197.

3. Colicino, E.G., and Hehnly, H. (2018). Regulating a key mitotic regulator, polo-like kinase 1 (PLK1). Cytoskeleton 75, 481–494.

4. Ohta, M., Zhao, Z., Wu, D., Wang, S., Harrison, J.L., Gómez-Cavazos, J.S., Desai, A., and Oegema, K.F. (2021). Polo-like kinase 1 independently controls microtubule-nucleating capacity and size of the centrosome. J. Cell Biol. 220.

5. Wueseke, O., Zwicker, D., Schwager, A., Wong, Y.L., Oegema, K., Jülicher, F., Hyman, A.A., and Woodruff, J.B. (2016). Polo-like kinase phosphorylation determines Caenorhabditis elegans centrosome size and density by biasing SPD-5 toward an assembly-competent conformation. Biol. Open 5, 1431–1440.

6. Lee, K., and Rhee, K. (2011). PLK1 phosphorylation of pericentrin initiates centrosome maturation at the onset of mitosis. J. Cell Biol. 195, 1093–1101.

7. Mittasch, M., Tran, V.M., Rios, M.U., Fritsch, A.W., Enos, S.J., Gomes, B.F., Bond, A., Kreysing, M., and Woodruff, J.B. (2020). Regulated changes in material properties underlie centrosome disassembly during mitotic exit. J. Cell Biol. 219.

8. Cabral, G., Laos, T., Dumont, J., and Dammermann, A. (2019). Differential Requirements for Centrioles in Mitotic Centrosome Growth and Maintenance. Dev. Cell 50, 355–366.e6.

9. Sepulveda, G., Antkowiak, M., Brust-Mascher, I., Mahe, K., Ou, T., Castro, N.M., Christensen, L.N., Cheung, L., Jiang, X., Yoon, D., et al. (2018). Co-translational protein targeting facilitates centrosomal recruitment of PCNT during centrosome maturation in vertebrates. Elife 7.

10. Rathbun, L.I., Aljiboury, A.A., Bai, X., Hall, N.A., Manikas, J., Amack, J.D., Bembenek, J.N., and Hehnly, H. (2020). PLK1- and PLK4-Mediated Asymmetric Mitotic Centrosome Size and Positioning in the Early Zebrafish Embryo. Curr. Biol. 30, 4519–4527.e3.

11. Gomez-Ferreria, M.A., Rath, U., Buster, D.W., Chanda, S.K., Caldwell, J.S., Rines, D.R., and Sharp, D.J. (2007). Human Cep192 Is Required for Mitotic Centrosome and Spindle Assembly. Curr. Biol. 17, 1960–1966.

12. Soung, N.-K., Kang, Y.H., Kim, K., Kamijo, K., Yoon, H., Seong, Y.-S., Kuo, Y.-L., Miki, T., Kim, S.R., Kuriyama, R., et al. (2006). Requirement of hCenexin for Proper Mitotic Functions of Polo-Like Kinase 1 at the Centrosomes. Mol. Cell. Biol. 26, 8316–8335.

13. Soung, N.K., Park, J.E., Yu, L.R., Lee, K.H., Lee, J.M., Bang, J.K., Veenstra, T.D., Rhee, K., and Lee, K.S. (2009). Plk1-Dependent and -Independent Roles of an ODF2 Splice Variant, hCenexin1, at the Centrosome of Somatic Cells. Dev. Cell 16, 539–550.

14. Chang, J., Seo, S.G., Lee, K.H., Nagashima, K., Bang, J.K., Kim, B.Y., Erikson, R.L., Lee, K.W., Lee, H.J., Park, J.E., et al. (2013). Essential role of Cenexin1, but not Odf2, in ciliogenesis. http://dx.doi.org/10.4161/cc.23585 12, 655–662.

15. Meng, L., Park, J.-E., Kim, T.-S., Lee, E.H., Park, S.-Y., Zhou, M., Bang, J.K., and Lee, K.S. (2015). Bimodal Interaction of Mammalian Polo-Like Kinase 1 and a Centrosomal Scaffold, Cep192, in the Regulation of Bipolar Spindle Formation. Mol. Cell. Biol. 35, 2626–2640.

16. Kemp, C.A., Kopish, K.R., Zipperlen, P., Ahringer, J., and O’Connell, K.F. (2004). Centrosome Maturation and Duplication in C. elegans Require the Coiled-Coil Protein SPD-2. Dev. Cell 6, 511–523.

17. Conduit, P.T., Richens, J.H., Wainman, A., Holder, J., Vicente, C.C., Pratt, M.B., Dix, C.I., Novak, Z.A., Dobbie, I.M., Schermelleh, L., et al. (2014). A molecular mechanism of mitotic centrosome assembly in Drosophila. Elife 3, 1–23.

18. Joukov, V., Walter, J.C., and De Nicolo, A. (2014). The Cep192-Organized Aurora A-Plk1 Cascade Is Essential for Centrosome Cycle and Bipolar Spindle Assembly. Mol. Cell 55, 578–591.

19. Colicino, E.G., Stevens, K., Curtis, E., Rathbun, L., Bates, M., Manikas, J., Amack, J.J., Freshour, J., and Hehnly, H. (2019). Chromosome misalignment is associated with PLK1 activity at cenexin-positive mitotic centrosomes. Mol. Biol. Cell 30, 1598–1609.

20. Ishikawa, H., Kubo, A., Tsukita, S., and Tsukita, S. (2005). Odf2-deficient mother centrioles lack distal/subdistal appendages and the ability to generate primary cilia. Nat. Cell Biol. 7, 517–524.

21. Tateishi, K., Yamazaki, Y., Nishida, T., Watanabe, S., Kunimoto, K., Ishikawa, H., and Tsukita, S. (2013). Two appendages homologous between basal bodies and centrioles are formed using distinct Odf2 domains. J. Cell Biol. 203, 417–425.

22. Hung, H.F., Hehnly, H., and Doxsey, S. (2016). The mother centriole appendage protein cenexin modulates lumen formation through spindle orientation. Curr. Biol. 26, 793–801.

23. Rusan, N.M., Serdar Tulu, U., Fagerstrom, C., and Wadsworth, P. (2002). Reorganization of the microtubule array in prophase/prometaphase requires cytoplasmic dynein-dependent microtubule transport. J. Cell Biol. 158, 997–1003.

24. Tulu, U.S., Rusan, N.M., and Wadsworth, P. (2003). Peripheral, Non-Centrosome-Associated Microtubules Contribute to Spindle Formation in Centrosome-Containing Cells. Curr. Biol. 13, 1894–1899.

25. Chen, C.T., Hehnly, H., Yu, Q., Farkas, D., Zheng, G., Redick, S.D., Hung, H.F., Samtani, R., Jurczyk, A., Akbarian, S., et al. (2014). A Unique Set of Centrosome Proteins Requires Pericentrin for Spindle-Pole Localization and Spindle Orientation. 24, 2327–2334.

26. Sahabandu, N., Kong, D., Magidson, V., Nanjundappa, R., Sullenberger, C., Mahjoub, M.R., and Loncarek, J. (2019). Expansion microscopy for the analysis of centrioles and cilia. J. Microsc. 276, 145–159.

27. Chozinski, T.J., Halpern, A.R., Okawa, H., Kim, H.-J., Tremel, G.J., Wong, R.O.L., and Vaughan, J.C. (2016). Expansion microscopy with conventional antibodies and fluorescent proteins. Nat. Methods 2016 136 13, 485–488.

28. Asano, S.M., Gao, R., Wassie, A.T., Tillberg, P.W., Chen, F., and Boyden, E.S. (2018). Expansion Microscopy: Protocols for Imaging Proteins and RNA in Cells and Tissues. Curr. Protoc. Cell Biol. 80, e56.

29. Levy, Y.Y., Lai, E.Y., Remillard, S.P., Heintzelman, M.B., and Fulton, C. (1996). Centrin Is a Conserved Protein That Forms Diverse Associations With Centrioles and MTOCs in Naegleria and Other Organisms. Cell Motil. Cytoskeleton 33, 298–323.

30. Salisbury, J.L., Suino, K.M., Busby, R., and Springett, M. (2002). Centrin-2 Is Required for Centriole Duplication in Mammalian Cells. Curr. Biol. 12, 1287–1292.

31. Wong, S.-S., Wilmott, Z.M., Saurya, S., Zhou, F.Y., Chau, K.-Y., Goriely, A., and Raff, J.W. (2021). Mother centrioles generate a local pulse of Polo/PLK1 activity to initiate mitotic centrosome assembly. bioRxiv, 2021.10.26.465695.

32. Novorol, C., Burkhardt, J., Wood, K.J., Iqbal, A., Roque, C., Coutts, N., Almeida, A.D., He, J., Wilkinson, C.J., and Harris, W.A. (2013). Microcephaly models in the developing zebrafish retinal neuroepithelium point to an underlying defect in metaphase progression. Open Biol. 3, 130065.

33. Aljiboury, A.A., Mujcic, A., Cammerino, T., Rathbun, L.I., and Hehnly, H. (2021). Imaging the early zebrafish embryo centrosomes following injection of small-molecule inhibitors to understand spindle formation. STAR Protoc. 2, 100293.

34. Colicino, E.G., Garrastegui, A.M., Freshour, J., Santra, P., Post, D.E., Kotula, L., and Hehnly, H. (2018). Gravin regulates centrosome function through PLK1. Mol. Biol. Cell 29.

35. Chozinski, T.J., Halpern, A.R., Okawa, H., Kim, H.J., Tremel, G.J., Wong, R.O.L., and Vaughan, J.C. (2016). Expansion microscopy with conventional antibodies and fluorescent proteins. Nat. Methods 13, 485–488.

36. Altschul, S.F., Gish, W., Miller, W., Myers, E.W., and Lipman, D.J. (1990). Basic local alignment search tool. J. Mol. Biol. 215, 403–410.

37. Edgar, R.C. (2004). MUSCLE: multiple sequence alignment with high accuracy and high throughput. Nucleic Acids Res. 32, 1792–1797.

38. Guindon, S., Dufayard, J.F., Lefort, V., Anisimova, M., Hordijk, W., and Gascuel, O. (2010). New Algorithms and Methods to Estimate Maximum-Likeli-hood Phylogenies: Assessing the Performance of PhyML 3.0. Syst. Biol. 59, 307–321.

39. Yu, G., Smith, D.K., Zhu, H., Guan, Y., and Lam, T.T.Y. (2017). ggtree: an r package for visualization and annotation of phylogenetic trees with their covariates and other associated data. Methods Ecol. Evol. 8, 28–36.

