## Supplemental figures and stats for "Pericentriolar matrix integrity relies on cenexin and Polo-Like Kinase (PLK)1"

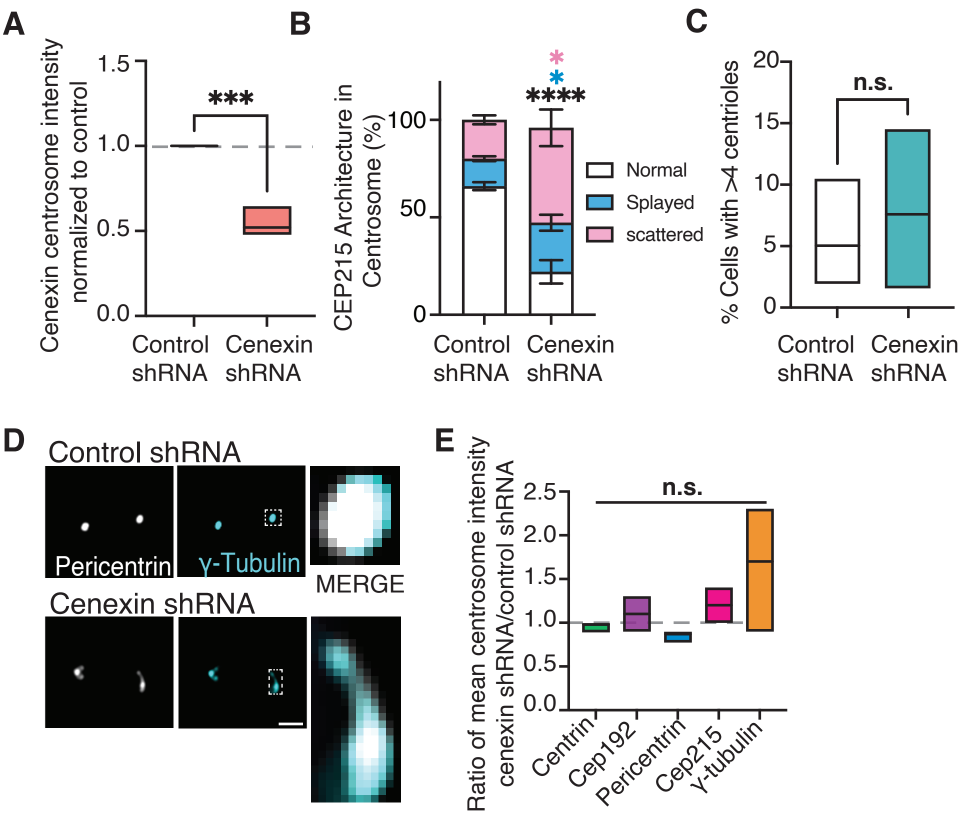


**Figure S1. Cenexin loss results in PCM specific fragmentation. Related to Figure 1.** (A) Box plot depicting cenexin centrosome intensity in mitotic control and cenexin shRNA normalized to control. Box boundaries denote the 25th and 75th percentiles. Unpaired, two-tailed Student's t-tests, ***p<0.001. (B) Stacked Bar graph depicting percentages of CEP215 architecture (scattered, purple; splayed, pink and normal, white) in control shRNA and cenexin shRNA treated cells. Means with SEM shown. Student t-test, *p<0.05, ***p<0.001. (C) Floating bars depicting percentage of control and cenexin shRNA cells with >4 centrioles. Min, max and median are displayed. Unpaired, two-tailed Student's t-tests, n.s. not significant. (D) Projections of control shRNA and cenexin shRNA immunolabeled for pericentrin (grey) and γ-tubulin (cyan). Insets at 3-4X magnification (merge). Scale bar, 5 µm. (E) Floating bars of the ration of mean centrosome intensity of centrosome marker proteins centrin (green), Cep192 (purple), Pericentrin (cyan), Cep215 (pink), and γ-tubulin (orange) in cenexin shRNA/control shRNA. Min, max and median are displayed. Grey dashed line represent ratio of 1. One-way ANOVA, n.s. not significant. For all graphs: detailed statistical analysis in Table S1.

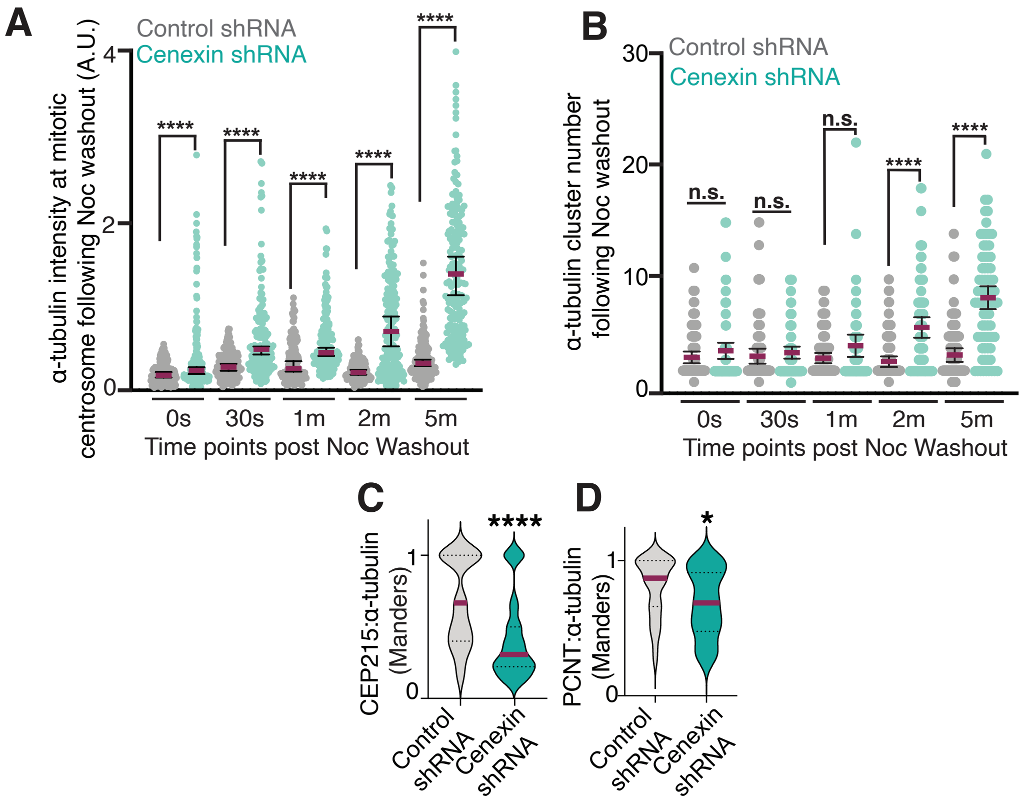


**Figure S2. Cenexin tempers microtubule nucleation by mediating pericentrin associated acentrosomal nucleation sites. Related to Figure 2.** (A-B) Scatter plot depicting α-tubulin intensity at mitotic centrosomes (A) and α-tubulin cluster number (B) 0s, 30s, 1m, 2m and 5m following nocodazole washout in control shRNA (grey) and cenexin shRNA (cyan) cells. Mean (magenta) with 95% confidence intervals are shown. Unpaired, two-tailed Student's t-tests performed between control and cenexin shRNA cells at each time point, n.s. not significant, ****$p<0.0001$. (C-D) Violin plot demonstrating colocalization of CEP215 (C) or pericentrin (D) with α-tubulin clusters in control shRNA (grey) and cenexin shRNA (cyan) treated cells measured using Mander's overlap coefficient. Magenta line denotes the median, dashed lines denote the 25th and 75th percentiles. Unpaired, two-tailed Student's t-tests, *$p<0.05$, ****$p<0.0001$. For all graphs: detailed statistical analysis in Table S1.

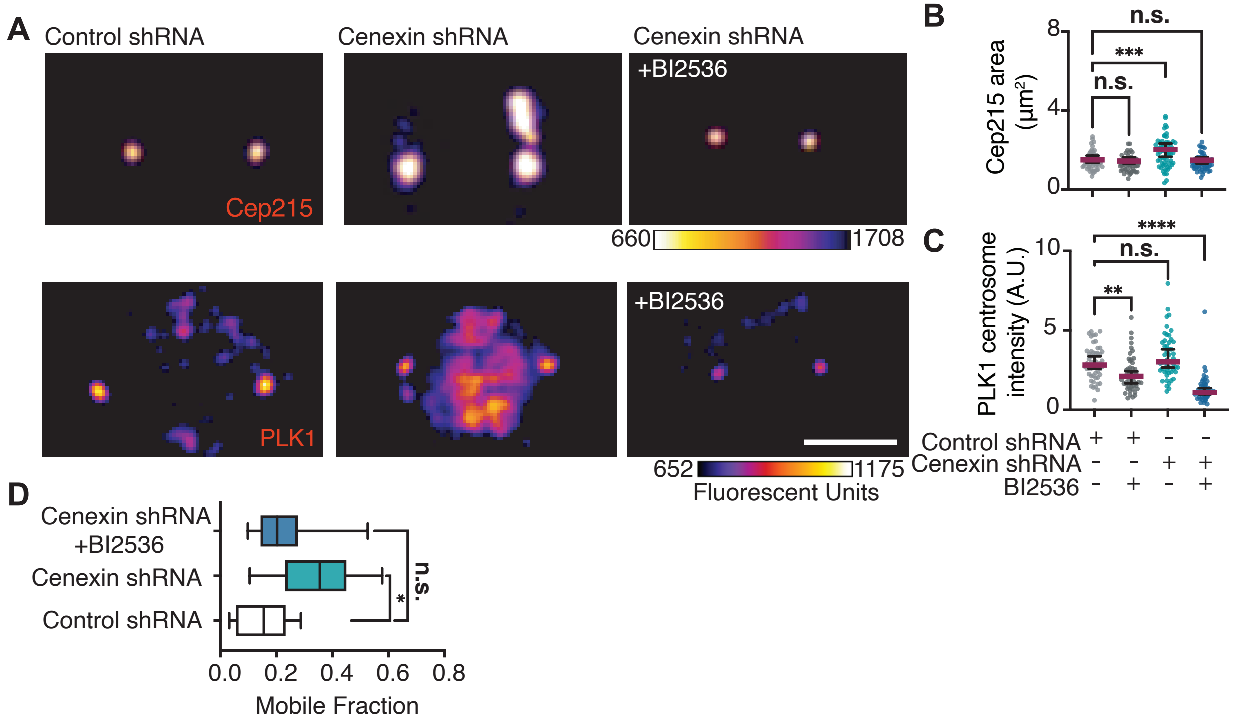


**Figure S3. Cenexin and PLK1 work together to maintain proper PCM organization. Related to Figure 3.** (A) Projections of mitotic control shRNA, cenexin shRNA and cenexin shRNA treated with BI2536 cells (Fire LUT) labeled for centrosome markers Cep215 (top) and PLK1 (bottom). Scale bar, 5 μm. (B-C) Scatter plots depicting Cep215 area ($\mu m^2$, B) and PLK1 mean centrosome intensity (A.U., C) under control shRNA and cenexin shRNA with and without BI2536. Mean (magenta) and 95% confidence intervals are shown. One-way ANOVA with multiple comparisons to control cells, n.s. not significant, $^{**}p<0.01$, $^{***}p<0.001$, $^{****}p<0.0001$. (D) Box-and-whisker plots of mobile fraction of RFP-PACT at metaphase spindle poles in control shRNA, cenexin shRNA and cenexin shRNA+BI2536. Box boundaries denote the 25th and 75th percentiles. One-way ANOVA with multiple comparisons to control cell, n.s. not significant, $^{*}p<0.05$. For all graphs: detailed statistical analysis in Table S1.

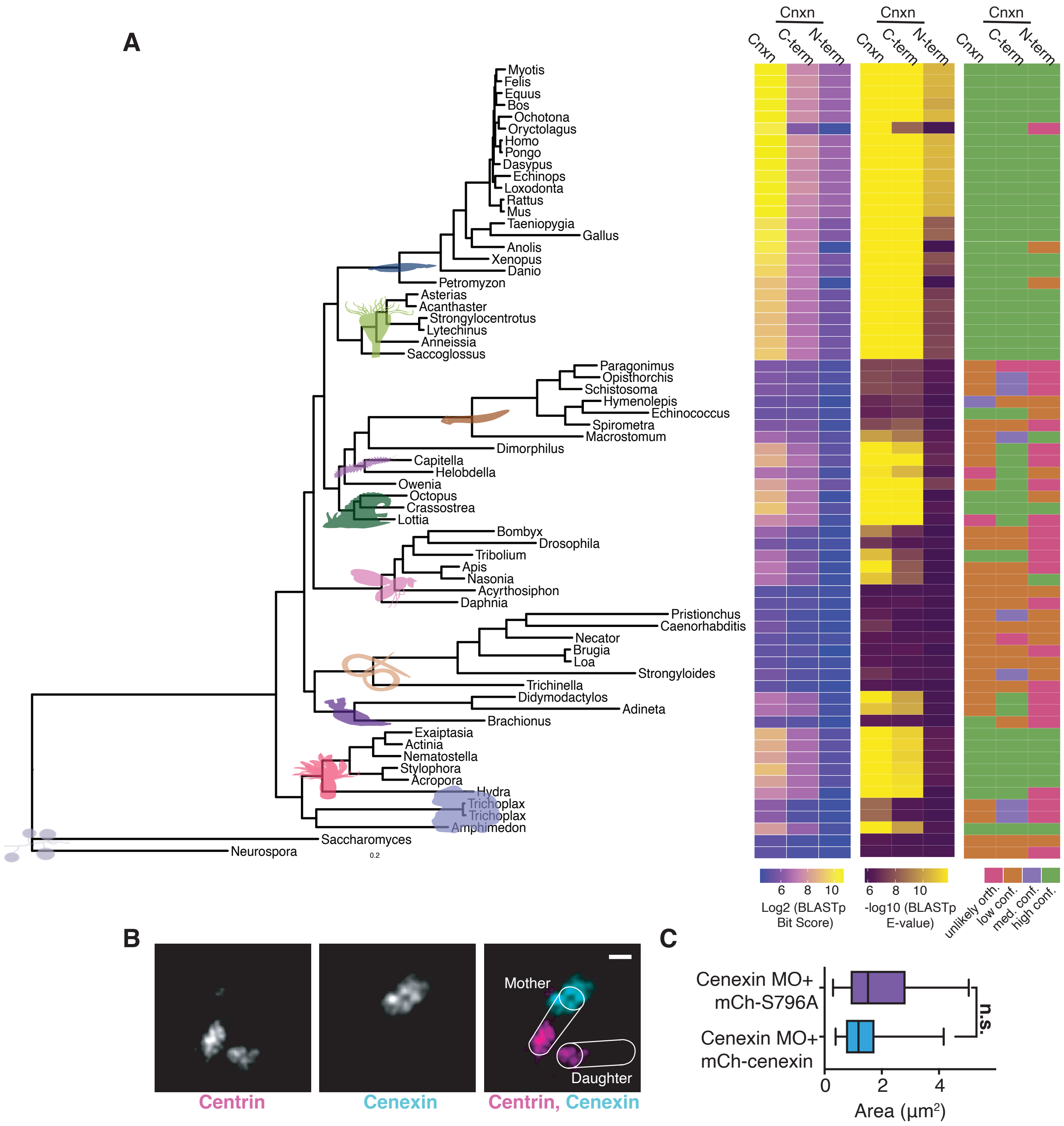


**Figure S4. Cenexin phosphorylation at its conserved C-terminal PLK1 binding site is required for maintenance of PCM in vivo. Related to Figure 4.** (A) BlastP searches using local BLAST instalation where hit confidence as potential orthologs were determined by hit length, percent identity, and NCBI annotation. High confidence orthologs were noted if NCBI annotated as a Cnxn ortholog or had a matched alignment length >100 a.a (Cnxn), >80 a.a (C-term), or >30 a.a. (N-term) and a percent identity >60% (Cnxn) or >40% (C-term). Additional considerations for unlikely, low confidence, and medium confidence discussed in methods. See Supplemental Files S1-S5. (B) Projections of expansion confocal images of an interphase Hela cell labeled for centrin (grey, magenta in merge) and cenexin (grey, cyan in merge). Centrioles are traced to show mother and daughter centrioles. Scale bar 1 µm. (C) Box-and-whisker plots of mCh-cenexin and mCh-cenexin-S796A area (µm$^2$) at mitotic centrosomes in cenexin depleted embryos (MO). Box boundaries denote the 25th and 75th percentiles. Unpaired, two-tailed Student's t-tests, n.s. not significant. For all graphs: detailed statistical analysis in Table S1.

| Figures | Category | n cells | n experiments | Statistical Test | Parameters | Result | p-value |
|---|---|---|---|---|---|---|---|
| 1B | Human cell (HeLa) control shRNA | 25 | n=3, representing 1 | Two-tailed Student's t-test | t=0.7746, df=98 | n.s. | 0.4404 |
| 1B | Human cell (HeLa) cenexin shRNA | 25 | n=3, representing 1 | Two-tailed Student's t-test | t=0.7746, df=98 | n.s. | 0.4404 |
| 1C | Human cell (HeLa) control shRNA | 25 | n=3, representing 1 | Two-tailed Student's t-test | t=0.3695, df=98 | n.s. | 0.7125 |
| 1C | Human cell (HeLa) cenexin shRNA | 25 | n=3, representing 1 | Two-tailed Student's t-test | t=0.3695, df=98 | n.s. | 0.7125 |
| 1D | Human cell (HeLa) control shRNA | 25 | n=3, representing 1 | Two-tailed Student's t-test | t=5.055, df=98 | **** | <0.0001 |
| 1D | Human cell (HeLa) cenexin shRNA | 25 | n=3, representing 1 | Two-tailed Student's t-test | t=5.055, df=98 | **** | <0.0001 |
| 1E | Human cell (HeLa) control shRNA | 25 | n=3, representing 1 | Two-tailed Student's t-test | t=6.469, df=106 | **** | <0.0001 |
| 1E | Human cell (HeLa) cenexin shRNA | 25 | n=3, representing 1 | Two-tailed Student's t-test | t=6.469, df=106 | **** | <0.0001 |
| 1F | Human cell (HeLa) control shRNA | 27 | n=3, representing 1 | Two-tailed Student's t-test | t=3.885, df=98 | *** | 0.0002 |
| 1F | Human cell (HeLa) cenexin shRNA | 25 | n=3, representing 1 | Two-tailed Student's t-test | t=3.885, df=98 | *** | 0.0002 |
| S1A | Human cell (HeLa) control shRNA | 75 | 3 | Two-tailed Student's t-test | t=8.953, df=4 | *** | 0.0009 |
| S1A | Human cell (HeLa) cenexin shRNA | 75 | 3 | Two-tailed Student's t-test | t=8.953, df=4 | *** | 0.0009 |
| S1B | Human cell (HeLa) control shRNA | >500 | 6 | Two-tailed Student's t-test | t=6.944, df=10 | Normal: **** | Normal: <0.0001 |
| S1B | Human cell (HeLa) control shRNA | >500 | 6 | Two-tailed Student's t-test | t=2.581, df=10 | Splayed: * | Splayed: 0.0273 |
| S1B | | >500 | 6 | Two-tailed Student's t-test | t=2.581, df=10 | Splayed: * | Splayed: 0.0273 |

| | | | | | | | |
|---|---|---|---|---|---|---|---|
| | Human cell (HeLa) cenexin shRNA | | | | t=2.988, df=10 | Scattered : * | Scattered: 0.0136 |
| S1C | Human cell (HeLa) control shRNA | | 6 | Two-tailed Student's t-test | t=1.136, df=9 | n.s. | 0.2854 |
| | Human cell (HeLa) cenexin shRNA | | 5 | | | | |
| S1E | CEP192 | >75 | 3 | One Way ANOVA | F (5, 12) = 2.399 | n.s. | P=0.0995 |
| | Pericentrin | | 3 | | | | |
| | CEP215 | | 3 | | | | |
| | γ-tubulin | | 3 | | | | |
| 2C | Human cell (HeLa) control shRNA | >174 | 3 | Two-tailed Student's t-test | t=17.30, df=359 | **** | <0.0001 |
| | Human cell (HeLa) cenexin shRNA | | 3 | | | | |
| 2D | Human cell (HeLa) control shRNA | >79 | 3 | Two-tailed Student's t-test | t=6.816, df=157 | **** | <0.0001 |
| | Human cell (HeLa) cenexin shRNA | | 3 | | | | |
| 2F | Human cell (HeLa) control shRNA | >76 | 3 | Two-tailed Student's t-test | t=4.281, df=162 | **** | <0.0001 |
| | Human cell (HeLa) cenexin shRNA | | 3 | | | | |
| 2G | Human cell (HeLa) control shRNA | >78 | 3 | Two-tailed Student's t-test | t=0.3306, df=164 | n.s. | 0.7414 |
| | Human cell (HeLa) cenexin shRNA | | 3 | | | | |
| S2A | Human cell (HeLa) | >158 | 3 | | t=5.107, df=321 | **** | <0.0001 |

| | | | | | | | |
|---|---|---|---|---|---|---|---|
| | control shRNA | | 3 | Two-tailed Student's t-test | t=7.961, df=337 | **** | |
| | Human cell (HeLa) cenexin shRNA | | 3 | | t=6.443, df=325 | **** | |
| | | | 3 | | t=11.73, df=347 | **** | |
| | | | 3 | | t=12.97, df=373 | **** | |
| S2B | Human cell (HeLa) control shRNA | >75 | 3 | Two-tailed Student's t-test | t=0.5556, df=153 | n.s. | 0.5793 |
| | | | 3 | | t=1.214, df=161 | n.s. | 0.2265 |
| | Human cell (HeLa) cenexin shRNA | | 3 | | t=0.5107, df=154 | n.s. | 0.6103 |
| | | | 3 | | t=4.333, df=157 | **** | <0.0001 |
| | | | 3 | | t=7.369, df=151 | **** | <0.0001 |
| S2C | Human cell (HeLa) control shRNA | >76 | n=3 | Two-tailed Student's t-test | t=5.516, df=152 | **** | <0.0001 |
| | Human cell (HeLa) cenexin shRNA | | n=3 | | | | |
| S2D | Human cell (HeLa) control shRNA | >27 | n=3, representing 1 | Two-tailed Student's t-test | | * | 0.0434 |
| | Human cell (HeLa) cenexin shRNA | | n=3, representing 1 | | | | |
| 3B | Human cell (HeLa) control shRNA | 25 | n=3, representing 1 | One Way ANOVA | F (3, 194) = 199.8 | Control | Control |
| | Human cell (HeLa) control shRNA+ BI2536 | 25 | n=3, representing 1 | | | **** | <0.0001 |
| | Human cell (HeLa) cenexin shRNA | 25 | n=3, representing 1 | | | **** | <0.0001 |
| | Human cell (HeLa) cenexin shRNA+ BI2536 | 25 | n=3, representing 1 | | | *** | <0.0002 |
| 3C | Human cell (HeLa) control shRNA | 25 | n=3, representing 1 | One Way ANOVA | F (3, 200) = 108.6 | Control | Control |

| | | | | | | | |
|---|---|---|---|---|---|---|---|
| | Human cell (HeLa) control shRNA+ BI2536 | 25 | n=3, representing 1 | | | **** | <0.0001 |
| | Human cell (HeLa) cenexin shRNA | 25 | n=3, representing 1 | | | **** | <0.0001 |
| | Human cell (HeLa) cenexin shRNA+ BI2536 | 25 | n=3, representing 1 | | | **** | <0.0001 |
| 3E | Human cell (HeLa) control shRNA | 7 | n=3, representing 1 | One Way ANOVA | F (3, 65) = 15.09 | Control | Control |
| | Human cell (HeLa) control shRNA+ BI2536 | 6 | n=3, representing 1 | | | n.s. | >0.9999 |
| | Human cell (HeLa) cenexin shRNA | 12 | n=3, representing 1 | | | **** | <0.0001 |
| | Human cell (HeLa) cenexin shRNA+ BI2536 | 9 | n=3, representing 1 | | | n.s. | 0.8196 |
| 3F | Human cell (HeLa) control shRNA | 13 | n=3, representing 1 | One Way ANOVA | F (3, 79) = 95.24 | Control | Control |
| | Human cell (HeLa) control shRNA+ BI2536 | 12 | n=3, representing 1 | | | * | 0.0163 |
| | Human cell (HeLa) cenexin shRNA | 8 | n=3, representing 1 | | | **** | <0.0001 |
| | Human cell (HeLa) cenexin shRNA+ BI2536 | 9 | n=3, representing 1 | | | n.s. | 0.2868 |
| 3I | Human cell (HeLa) control shRNA | 6 | >3 | | | | |
| | Human cell (HeLa) cenexin shRNA | 8 | | | | | |

| | | | | | | | |
|---|---|---|---|---|---|---|---|
| | Human cell (HeLa) cenexin shRNA+ BI2536 | 10 | | | | | |
| S3B | Human cell (HeLa) control shRNA | 25 | n=3, representing 1 | One Way ANOVA | F (3, 215) = 13.32 | Control | Control |
| | Human cell (HeLa) control shRNA+ BI2536 | 27 | n=3, representing 1 | | | n.s. | 0.4275 |
| | Human cell (HeLa) cenexin shRNA | 29 | n=3, representing 1 | | | *** | 0.0001 |
| | Human cell (HeLa) cenexin shRNA+ BI2536 | 27 | n=3, representing 1 | | | n.s. | 0.7262 |
| S3C | Human cell (HeLa) control shRNA | 26 | n=3, representing 1 | One Way ANOVA | F (3, 195) = 30.64 | Control | Control |
| | Human cell (HeLa) control shRNA+ BI2536 | 26 | n=3, representing 1 | | | ** | 0.0083 |
| | Human cell (HeLa) cenexin shRNA | 25 | n=3, representing 1 | | | n.s. | 0.1330 |
| | Human cell (HeLa) cenexin shRNA+ BI2536 | 25 | n=3, representing 1 | | | **** | <0.0001 |
| S3D | Human cell (HeLa) control shRNA | 6 | >3 | One Way ANOVA | F (2, 22) = 4.693 | Control | Control |
| | Human cell (HeLa) cenexin shRNA | 8 | | | | * | 0.0131 |
| | Human cell (HeLa) cenexin shRNA+ BI2536 | 10 | | | | n.s. | 0.3848 |
| 4F | Control injection | 12, 3 embryos | 1 | One Way ANOVA | F (3, 82) = 28.98 | Control | Control |

| | | | | | | | |
|---|---|---|---|---|---|---|---|
| | Cenexin MO injection | 13, 3 embryos | 1 | | | ** | 0.0092 |
| | Cenexin MO injection +mCherry cenexin-WT | 15, 4 embryos | 1 | | | n.s. | 0.3406 |
| | Cenexin MO injection +mCherry cenexin-S796A | 12, 4 embryos | 1 | | | **** | <0.0001 |
| 4G | Cenexin MO injection +mCherry cenexin-WT | 14, 3 embryos | 1 | Two-tailed Student's t-test | t=4.348, df=50 | **** | <0.0001 |
| | Cenexin MO injection +mCherry cenexin-S796A | 13, 3 embryos | 1 | | | | |
| S4B | Cenexin area ($\mu m^2$) | 10, 3 embryos | 1 | Two-tailed Student's t-test | t=1.715, df=34 | n.s. | 0.0955 |
| | | 8, 3 embryos | 1 | | | | |

**Table S1. Detailed statistical analysis of results reported in this study. Related to STAR methods.**
