## Supplemental File SF1 for "Pericentriolar matrix integrity relies on cenexin and Polo-Like Kinase (PLK)1": SF1C_phylo_analysis.html


### Phylogenetic analysis of Cennexin¶

Load the required libraries:

In [2]:

```
library("EBImage")
library("ggimage")
library("ggtree")
library("ggtreeExtra")
library("magrittr")
library("msa")
library("phytools")
library("rphylopic")
library("tidytree")
library("tidyverse")
library("treeio")
```

Load the raw tree in Newick format:

In [3]:

```
Ntree <- read.tree(file = "FINAL_SETS/nuclear_genes_tree.nwk")
Ntree
```

```
Phylogenetic tree with 67 tips and 65 internal nodes.

Tip labels:
  Loa_loa, Brugia_malayi, Necator_americanus, Caenorhabditis_elegans, Pristionchus_pacificus, Strongyloides_ratti, ...
Node labels:
  , 1.000000, 1.000000, 0.999998, 1.000000, 1.000000, ...

Unrooted; includes branch lengths.
```

Examine the tree to identify the outgroup node:

In [4]:

```
options(repr.plot.width = 22, repr.plot.height = 20)
ggtree(Ntree) + geom_tiplab(size = 7) + geom_label2(aes(label=node), size=3)
```

Use the outgroup node to reroot the tree

In [5]:

```
newTree <- reroot(tree = Ntree, node.number = 79)
```

Identify all the major phyla and annotate the tree with phylum identity:

In [6]:

```
options(repr.plot.width = 30, repr.plot.height = 20)
ggtree(newTree, branch.length = FALSE) + 
    geom_tiplab(size = 7) +
    geom_strip('Owenia_fusiformis', 'Dimorphilus_gyrociliatus', barsize=3, label="Annelida", offset=0.6, fontsize = 14, offset.text = 0.03) +
    geom_cladelab(node=81, label="Arthropoda", offset=0.6, barsize=2, align = TRUE, fontsize = 14) +
    geom_strip('Neurospora_crassa', 'Saccharomyces_cerevisiae', barsize=3, label="Ascomycota", offset=.6, fontsize = 14, offset.text = 0.03) +
    geom_cladelab(node=107, label="Chordata", offset=.6, barsize=2, align = TRUE, fontsize = 14) +
    geom_cladelab(node=126, label="Cnidaria", offset=.6, barsize=2, align = TRUE, fontsize = 14) +
    geom_cladelab(node=103, label="Echinodermata", offset=.6, barsize=2, align = TRUE, fontsize = 14) +
    geom_cladelab(node=41, label="Hemichordata", offset=.6, barsize=2, align = TRUE, fontsize = 14) +
    geom_cladelab(node=99, label="Mollusca", offset=.6, barsize=2, align = TRUE, fontsize = 14) +
    geom_cladelab(node=72, label="Nematoda", offset=.6, barsize=2, align = TRUE, fontsize = 14) +
    geom_cladelab(node=132, label="Placozoa", offset=.6, barsize=2, align = TRUE, fontsize = 14) +
    geom_cladelab(node=93, label="Platyhelminthes", offset=.6, barsize=2, align = TRUE, fontsize = 14) +
    geom_cladelab(node=17, label="Porifera", offset=.6, barsize=2, align = TRUE, fontsize = 14) +
    geom_cladelab(node=78, label="Rotifera", offset=.6, barsize=2, align = TRUE, fontsize = 14) +
    ggplot2::xlim(0, 4)
ggsave("base_tree.pdf", width = 30, height = 20)
```

Load the taxon annotation file:

In [7]:

```
anotInfo <- read.csv("FINAL_SETS/TAXONS_TO_USE_TABBLE_UPDATED.csv", sep = ",") %>% mutate(species = gsub(" ", "_", Taxon))
head(anotInfo)
```

A data.frame: 6 × 13

|  | code | Taxon | Taxon\_id | Genus | Family | Order | Class | Phylum | Kingdom | Superkingdom | protein\_fasta | misc | species |
| --- | --- | --- | --- | --- | --- | --- | --- | --- | --- | --- | --- | --- | --- |
|  | <chr> | <chr> | <chr> | <chr> | <chr> | <chr> | <chr> | <chr> | <chr> | <chr> | <chr> | <chr> | <chr> |
| 1 | hx | Helobdella robusta | Helobdella robusta | Helobdella | Glossiphoniidae | Rhynchobdellida | Clitellata | Annelida | Metazoa | Eukaryota | https://ftp.ncbi.nlm.nih.gov/genomes/all/GCF/000/326/865/GCF\_000326865.1\_Helobdella\_robusta\_v1.0/GCF\_000326865.1\_Helobdella\_robusta\_v1.0\_protein.faa.gz |  | Helobdella\_robusta |
| 2 | i1 | Capitella teleta | Capitella¬†sp. I¬† | Capitella | Capitellidae | Capitellida | Polychaeta | Annelida | Metazoa | Eukaryota | https://ftp.ncbi.nlm.nih.gov/genomes/all/GCA/000/328/365/GCA\_000328365.1\_Capca1/GCA\_000328365.1\_Capca1\_protein.faa.gz | capitella teleta | Capitella\_teleta |
| 3 |  | Owenia fusiformis |  | Owenia | Oweniidae | Sabellida | Polychaeta | Annelida | Metazoa | Eukaryota | https://ftp.ncbi.nlm.nih.gov/genomes/all/GCA/903/813/345/GCA\_903813345.1\_Owenia\_assembly\_annotated/GCA\_903813345.1\_Owenia\_assembly\_annotated\_protein.faa.gz |  | Owenia\_fusiformis |
| 4 |  | Dimorphilus gyrociliatus |  | Dimorphilus | Dinophilidae |  | Polychaeta | Annelida | Metazoa | Eukaryota | https://ftp.ncbi.nlm.nih.gov/genomes/all/GCA/904/063/045/GCA\_904063045.1\_Dgyrociliatus\_assembly/GCA\_904063045.1\_Dgyrociliatus\_assembly\_protein.faa.gz |  | Dimorphilus\_gyrociliatus |
| 5 | nz | Nasonia vitripennis | Nasonia vitripennis | Nasonia | Pteromalidae | Hymenoptera | Insecta | Arthropoda | Metazoa | Eukaryota | https://ftp.ncbi.nlm.nih.gov/genomes/all/GCF/009/193/385/GCF\_009193385.2\_Nvit\_psr\_1.1/GCF\_009193385.2\_Nvit\_psr\_1.1\_protein.faa.gz |  | Nasonia\_vitripennis |
| 6 | ai | Apis mellifera | Apis mellifera¬†38.2d | Apis | Apidae | Hymenoptera | Insecta | Arthropoda | Metazoa | Eukaryota | https://ftp.ncbi.nlm.nih.gov/genomes/all/GCF/003/254/395/GCF\_003254395.2\_Amel\_HAv3.1/GCF\_003254395.2\_Amel\_HAv3.1\_protein.faa.gz |  | Apis\_mellifera |

Rename the tips as genera only:

In [8]:

```
anotInfo %>% select(Taxon, Genus, Family, Order, Class, Phylum) %>% arrange(Taxon) -> phylo_info
d <- data.frame(label = sort(newTree$tip.label), phylo_info) 
tr2 = rename_taxa(newTree, d, label, Genus)
```

Load the home-brew BLAST results tables,

In [9]:

```
odf2_Nterm_blast <- read.csv("BLAST_results/odf2_NtermExt_blast_results.outfmt6", sep = "\t", header = T) %>% mutate(species = gsub("Trichoplax_sp1", "TrichoplaxSp1_", species)) %>% separate(species, into = c("Genus", "Species"))
odf2_Cterm_blast <- read.csv("BLAST_results/odf2_Cterm_blast_results.outfmt6", sep = "\t", header = T) %>% mutate(species = gsub("Trichoplax_sp1", "TrichoplaxSp1_", species)) %>% separate(species, into = c("Genus", "Species"))
odf2_iso9_blast <- read.csv("BLAST_results/odf2_iso9_blast_results.outfmt6", sep = "\t", header = T) %>% mutate(species = gsub("Trichoplax_sp1", "TrichoplaxSp1_", species)) %>% separate(species, into = c("Genus", "Species"))
IFTs_blast <- read.csv("BLAST_results/IFT_blast_results.outfmt6", sep = "\t", header = T)%>% mutate(species = gsub("Trichoplax_sp1", "TrichoplaxSp1_", species)) %>% separate(species, into = c("Genus", "Species"))
```

```
Warning message:
“Expected 2 pieces. Additional pieces discarded in 2 rows [11, 61].”
Warning message:
“Expected 2 pieces. Additional pieces discarded in 2 rows [12, 64].”
Warning message:
“Expected 2 pieces. Additional pieces discarded in 2 rows [14, 77].”
Warning message:
“Expected 2 pieces. Additional pieces discarded in 279 rows [1579, 1580, 1581, 1582, 1583, 1584, 1585, 1586, 1587, 1588, 1589, 1590, 1591, 1592, 1593, 1594, 1595, 1596, 1597, 1598, ...].”
```

Load the protein info for each BLAST table:

In [10]:

```
odf2_Nterm_info <- read.csv("BLAST_results/odf2_NtermExt_blast_results_protein_info.txt", sep = "\t", header =F) %>% rename(species = V1, protein_id = V2, description = V3) %>% mutate(species = gsub("Trichoplax sp", "TrichoplaxSp1", species)) %>% separate(species, into = c("Genus", "Species"))
odf2_Cterm_info <- read.csv("BLAST_results/odf2_CtermExt_blast_results_protein_info.txt", sep = "\t", header = F) %>% rename(species = V1, protein_id = V2, description = V3) %>% mutate(species = gsub("Trichoplax sp", "TrichoplaxSp1", species)) %>% separate(species, into = c("Genus", "Species"))
odf2_iso9_info <- read.csv("BLAST_results/odf2_iso9_blast_results_protein_info.txt", sep = "\t", header = F) %>% rename(species = V1, protein_id = V2, description = V3) %>% mutate(species = gsub("Trichoplax sp", "TrichoplaxSp1", species)) %>% separate(species, into = c("Genus", "Species"))
IFTs_info <- read.csv("BLAST_results/IFT_blast_results_protein_info.txt", sep = "\t", header = F) %>% rename(species = V1, protein_id = V2, description = V3) %>% mutate(species = gsub("Trichoplax sp", "TrichoplaxSp1", species)) %>% separate(species, into = c("Genus", "Species"))
```

```
Warning message:
“Expected 2 pieces. Additional pieces discarded in 2 rows [44, 55].”
Warning message:
“Expected 2 pieces. Additional pieces discarded in 2 rows [44, 55].”
Warning message:
“Expected 2 pieces. Additional pieces discarded in 2 rows [44, 55].”
Warning message:
“Expected 2 pieces. Additional pieces discarded in 46 rows [643, 644, 645, 646, 647, 648, 649, 650, 651, 652, 653, 654, 655, 656, 657, 658, 659, 660, 661, 662, ...].”
```

Load the NCBI BLAST results and descriptions tables:

In [11]:

```
odf2_Cterm_ncbi_descr <- read.csv("NCBI_BLAST_results/odf2_Cterm/XEA3CW8P013-Alignment-Descriptions.csv", header = T, sep = ",") %>% 
        mutate(Accession = gsub('.*","', '', Accession), Accession = gsub('")', '', Accession), Genus = gsub(" .*", "", Scientific.Name	))

odf2_Nterm_ncbi_descr <- read.csv("NCBI_BLAST_results/odf2_Nterm/XE9YX3TG016-Alignment-Descriptions.csv", header = T, sep = ",") %>% 
        mutate(Accession = gsub('.*","', '', Accession), Accession = gsub('")', '', Accession), Genus = gsub(" .*", "", Scientific.Name	)) 

odf2_iso9_ncbi_descr <- read.csv("NCBI_BLAST_results/odf2_iso9/XE9H7JZF013-Alignment-Descriptions.csv", header = T, sep = ",") %>% 
        mutate(Accession = gsub('.*","', '', Accession), Accession = gsub('")', '', Accession), Genus = gsub(" .*", "", Scientific.Name	))

CEP192_ncbi_descr <- read.csv("NCBI_BLAST_results/CEP192/XEAGKX23016-Alignment-Descriptions.csv", header = T, sep = ",") %>% 
        mutate(Accession = gsub('.*","', '', Accession), Accession = gsub('")', '', Accession), Genus = gsub(" .*", "", Scientific.Name	))

Centrin_ncbi_descr <- read.csv("NCBI_BLAST_results/Centrin/XEASXN46016-Alignment-Descriptions.csv", header = T, sep = ",") %>% 
        mutate(Accession = gsub('.*","', '', Accession), Accession = gsub('")', '', Accession), Genus = gsub(" .*", "", Scientific.Name	))
```

Add phyla images to tree:

In [12]:

```
options(repr.plot.width = 10, repr.plot.height = 10)
p1 <- ggtree(tr2) + 
    geom_tiplab(size = 7, offset = 3) +
    geom_strip('Owenia', 'Dimorphilus', barsize=3, label="Annelida", offset=0.6, fontsize = 14, offset.text = 0.03) +
    geom_cladelab(node=81, label="Arthropoda", offset=0.6, barsize=2, align = TRUE, fontsize = 14) +
    geom_strip('Neurospora', 'Saccharomyces', barsize=3, label="Ascomycota", offset=.6, fontsize = 14, offset.text = 0.03) +
    geom_cladelab(node=107, label="Chordata", offset=.6, barsize=2, align = TRUE, fontsize = 14) +
    geom_cladelab(node=126, label="Cnidaria", offset=.6, barsize=2, align = TRUE, fontsize = 14) +
    geom_cladelab(node=103, label="Echinodermata", offset=.6, barsize=2, align = TRUE, fontsize = 14) +
    geom_cladelab(node=41, label="Hemichordata", offset=.6, barsize=2, align = TRUE, fontsize = 14) +
    geom_cladelab(node=99, label="Mollusca", offset=.6, barsize=2, align = TRUE, fontsize = 14) +
    geom_cladelab(node=72, label="Nematoda", offset=.6, barsize=2, align = TRUE, fontsize = 14) +
    geom_cladelab(node=132, label="Placozoa", offset=.6, barsize=2, align = TRUE, fontsize = 14) +
    geom_cladelab(node=93, label="Platyhelminthes", offset=.6, barsize=2, align = TRUE, fontsize = 14) +
    geom_cladelab(node=17, label="Porifera", offset=.6, barsize=2, align = TRUE, fontsize = 14) +
    geom_cladelab(node=78, label="Rotifera", offset=.6, barsize=2, align = TRUE, fontsize = 14) +
    geom_treescale() +
    ggplot2::xlim(0, 3)
```

We can include the phylopic images to the tree if we know the requisite nodes:

In [13]:

```
phylopic_info <- data.frame(node = c(107, 41, 81, 72, 99, 91, 93, 126, 17, 132, 133, 103, 78),
                            phylopic = c("aa9d2eb5-d86b-4fe5-adee-99056db1d8d8", 
                                        "5f5db6a9-5ca9-4297-b3ab-82a7a42627ba", 
                                        "48bcbd0d-a902-4be8-9acc-60a6fbfd5f8d", 
                                        "e9bf2dfa-42fe-4074-adad-e7a2a4be21e9",
                                        "ebf8143a-5bba-4d0f-9897-e8e752d11e37", 
                                        "e7d59ad8-887c-4017-bad5-6e2b5b167dc6", 
                                        "f87cf0c9-dd85-4165-8936-0136b971891b", 
                                        "d148ee59-7247-4d2a-a62f-77be38ebb1c7", 
                                        "3449d9ef-2900-4309-bf22-5262c909344b",
                                        "9331b2ea-bb3f-493e-af00-c4a4baa5a1f8",
                                        "31aa0781-ca28-4362-8955-520f1c45f232",
                                        "4a56d23f-0725-4a13-bda7-665e0d5cf9b8",
                                        "3042a73d-353d-4191-811f-9b12f57c958c"),
                            phylum = c("Chordates", "Hemichordates", "Arthropods", 
                                       "Nematodes", "Molluscs", "Annelids",
                                      "Platyhelminthes", "Cnidaria", "Porifera",
                                      "Placozoa", "Ascomycota (Fungi)", "Echinoderms", "Rotifers"))
p3 <- p1 %<+% phylopic_info + geom_nodelab(aes(image=phylopic, colour = phylum), geom="phylopic", alpha=.8) + theme(legend.position = "none")
```

###### The nodes represented the major phyla are as follows:¶

Annelida = paraphyletic('Owenia\_fusiformis', 'Dimorphilus\_gyrociliatus')

Arthropoda = 81

Ascomycota = ('Neurospora\_crassa', 'Saccharomyces\_cerevisiae')

Chordata = 107

Cnidaria = 126

Echinodermata = 103

Hemichordata = 41

Mollusca = 99

Nematoda = 72

Placozoa = 132

Platyhelminthes = 93

Porifera = 17

Rotifera = 78

Here's an example tree with phylopic images:

In [19]:

```
options(repr.plot.width = 10, repr.plot.height = 5)
ggtree(tr2) + geom_cladelab(data = phylopic_info, 
                mapping = aes(node = node, label = phylum, image = phylopic, colour = phylum), 
                geom = "phylopic")  %>% suppressWarnings()
```

Let's create *ad hoc* categories of BLAST hits using our homebrew BLASTp resutls:

In [20]:

```
merge(odf2_iso9_blast, odf2_iso9_info, by.x = "saccver", by.y = "protein_id")%>% 
    group_by(Genus.x) %>% 
    slice(which.min(evalue)) %>% 
    select(Genus.x, pident, evalue, length, mismatch, bitscore, description)  %>% 
    mutate(odf2_status = ifelse(grepl("outer dense fiber", description), 
                                "high conf.", 
                                ifelse(length > 100 & pident > 60 & !grepl("outer dense fiber", description), 
                                       "high conf.", 
                                       ifelse(length < 100  & length > 30 & pident > 40 & !grepl("outer dense fiber", description), 
                                              "medium conf.", 
                                              ifelse(length > 30 & pident < 40 & !grepl("outer dense fiber", description), 
                                                     "low conf.", 
                                                     "unlikely orth.")))),
          iso9_bitscore = log2(bitscore),
          iso9_evalue = -log10(evalue+1E-30)) %>% 
    select(Genus.x, odf2_status, iso9_bitscore, iso9_evalue) -> odf2_status

merge(odf2_Cterm_blast, odf2_Cterm_info, by.x = "saccver", by.y = "protein_id")%>% 
    group_by(Genus.x) %>% 
    slice(which.min(evalue)) %>% 
    select(Genus.x, pident, evalue, length, mismatch, description, bitscore)  %>% 
    mutate(Cterm_status = ifelse(grepl("outer dense fiber", description), 
                                "high conf.", 
                                ifelse(length > 80 & pident > 40 & !grepl("outer dense fiber", description), 
                                       "high conf.", 
                                       ifelse(length < 80  & length > 30 & pident > 40 & !grepl("outer dense fiber", description), 
                                              "medium conf.", 
                                              ifelse(length > 30 & pident < 40 & !grepl("outer dense fiber", description), 
                                                     "low conf.", 
                                                     "unlikely orth.")))),
          Cterm_bitscore = log2(bitscore),
          Cterm_evalue = -log10(evalue+1E-30)) %>% 
    select(Genus.x, Cterm_status, Cterm_bitscore, Cterm_evalue) -> Cterm_status

merge(odf2_Nterm_blast, odf2_Nterm_info, by.x = "saccver", by.y = "protein_id")%>% 
    group_by(Genus.x) %>% 
    slice(which.min(evalue)) %>% 
    select(Genus.x, pident, evalue, length, mismatch, description, bitscore)  %>% 
    mutate(Nterm_status = ifelse(grepl("outer dense fiber", description), 
                                "high conf.", 
                                ifelse(length > 30 & pident > 40 & !grepl("outer dense fiber", description), 
                                       "high conf.", 
                                       ifelse(length < 40  & length > 30 & pident > 40 & !grepl("outer dense fiber", description), 
                                              "medium conf.", 
                                              ifelse(length > 15 & pident < 40 & !grepl("outer dense fiber", description), 
                                                     "low conf.", 
                                                     "unlikely orth.")))),
          Nterm_bitscore = log2(bitscore),
          Nterm_evalue = -log10(evalue+1E-30)) %>% 
    select(Genus.x, Nterm_status, Nterm_bitscore, Nterm_evalue) -> Nterm_status
```

Combine these accordingly:

In [21]:

```
cbind(Cterm_status, odf2_status, Nterm_status) %>% select(Genus.x...1, iso9_bitscore, Cterm_bitscore, Nterm_bitscore) %>% as.data.frame %>% column_to_rownames(var = "Genus.x...1") -> odf2_bitscore_table
cbind(Cterm_status, odf2_status, Nterm_status) %>% select(Genus.x...1, iso9_evalue, Cterm_evalue, Nterm_evalue) %>% as.data.frame %>% column_to_rownames(var = "Genus.x...1") -> odf2_evalue_table
cbind(Cterm_status, odf2_status, Nterm_status) %>% select(Genus.x...1, odf2_status, Cterm_status, Nterm_status) %>% as.data.frame %>% column_to_rownames(var = "Genus.x...1") -> odf2_status_table
```

```
New names:
* Genus.x -> Genus.x...1
* Genus.x -> Genus.x...5
* Genus.x -> Genus.x...9

New names:
* Genus.x -> Genus.x...1
* Genus.x -> Genus.x...5
* Genus.x -> Genus.x...9

New names:
* Genus.x -> Genus.x...1
* Genus.x -> Genus.x...5
* Genus.x -> Genus.x...9
```

Also create an object with detection information from the NCBI blast (we'll add the human ones manually because we didn;t include humans as one of the target species... duh!):

In [22]:

```
merge(odf2_iso9_blast, odf2_iso9_info, by.x = "saccver", by.y = "protein_id")%>% 
    group_by(Genus.x) %>% 
    slice(which.min(evalue)) %>% 
    select(Genus.x, evalue, length, mismatch, description, bitscore)  %>% 
    mutate(odf2_iso9 = ifelse(Genus.x %in% odf2_iso9_ncbi_descr$Genus, "detected", "not detected"),
           odf2_Nterm = ifelse(Genus.x %in% odf2_Nterm_ncbi_descr$Genus, "detected", "not detected"),
           odf2_Cterm = ifelse(Genus.x %in% odf2_Cterm_ncbi_descr$Genus, "detected", "not detected"),
          CEP192 = ifelse(Genus.x %in% CEP192_ncbi_descr$Genus, "detected", "not detected"),
          Centrin = ifelse(Genus.x %in% Centrin_ncbi_descr$Genus, "detected", "not detected")) %>%
    mutate(odf2_iso9 = ifelse(Genus.x == "Homo", "detected", odf2_iso9),
           odf2_Nterm = ifelse(Genus.x == "Homo", "detected", odf2_Nterm),
           odf2_Cterm = ifelse(Genus.x == "Homo", "detected", odf2_Cterm),
          CEP192 = ifelse(Genus.x == "Homo", "detected", CEP192),
          Centrin = ifelse(Genus.x == "Homo", "detected", Centrin)) %>% 
    select(Genus.x, odf2_iso9, odf2_Cterm, odf2_Nterm, CEP192, Centrin) %>% 
    column_to_rownames(var = "Genus.x")-> All_status
```

For the main figure, let's annotate the tree with the NCBI hit resutls:

In [28]:

```
options(repr.plot.width = 8, repr.plot.height = 7)
p4 <- ggtree(tr2) + geom_cladelab(data = phylopic_info, offset.text = -0.2, align = T, offset = 0.2,imagesize = 0.04,
                mapping = aes(node = node, label = phylum, image = phylopic, colour = phylum), 
                geom = "phylopic") + 
            scale_colour_manual(values = c("#003b76","#92b74f","#0038d6","#9e5000","#cb54ff",
                                           "#005826","#ff6fe6","#d89f7b","#5a0091","#ff446f",
                                           "#868aff","#3c004c","#aca8c5"),
                                breaks = c('Chordates','Echinoderms','Hemichordates','Platyhelminthes',
                                           'Annelids','Molluscs','Arthropods','Nematodes','Rotifers',
                                           'Cnidaria','Placozoa','Porifera','Ascomycota (Fungi)')) +
            geom_hilight(node=61, fill="steelblue", alpha=1) +
            geom_hilight(node=50, fill="pink", alpha=1) 
gheatmap(p4, 
         All_status, 
         colnames_position = "top", 
         width = 0.55, 
         offset=0.25, 
         hjust = 0,
         colnames_angle = 35, 
         legend_title = "BLASTp\nresult", 
         colnames_offset_y = 0,
         font.size = 3,
         custom_column_labels = c("odf2 iso9", "odf2 C-term", "odf2 N-term", "Cep192", "Centrin"),
        ) + 
        scale_fill_manual(values = c("#009ad6","#a64200")) + 
        ggtree::vexpand(.4, -1) +
        scale_x_ggtree() + 
        ggplot2::ylim(-0.5, NA) +
        coord_cartesian(clip = "off") +
        guides(fill=guide_legend(title="BLASTp\nresult"))
# p5 
ggsave("Cennexin_and_co_phylogeny_v3.pdf", width = 8, height = 7)
```

```
Scale for 'fill' is already present. Adding another scale for 'fill', which
will replace the existing scale.

Scale for 'y' is already present. Adding another scale for 'y', which will
replace the existing scale.
```

For the homebrew BLASTp, let's create a tree with each of the three heatmap results summarizing the BLASTp info. First, using the Bit scores:

In [31]:

```
p4 %<+% phylopic_info + geom_nodelab(aes(image=phylopic, colour = phylum), geom="phylopic", alpha=.7) + theme(legend.position = "none")+ 
            scale_colour_manual(values = c("#003b76","#92b74f","#0038d6","#9e5000","#cb54ff",
                                           "#005826","#ff6fe6","#d89f7b","#5a0091","#ff446f",
                                           "#868aff","#3c004c","#aca8c5"),
                                breaks = c('Chordates','Echinoderms','Hemichordates','Platyhelminthes',
                                           'Annelids','Molluscs','Arthropods','Nematodes','Rotifers',
                                           'Cnidaria','Placozoa','Porifera','Ascomycota (Fungi)')) -> p10
options(repr.plot.width = 20, repr.plot.height = 20)
gheatmap(p10, low = "blue", high = "yellow",font.size = 8,
         odf2_bitscore_table, 
         colnames_position = "top", 
         width = 0.3, 
         offset=0.3,
         colnames_angle = 0, 
         legend_title = "log2(BLASTp\nBit Score)", 
         colnames_offset_y = 1, 
         custom_column_labels = c("Cnxn", "Cnxn\nC-ter", "Cnxn\nN-ter")) + ggplot2::ylim(-0.5, NA) + theme(legend.position="bottom")
ggsave("supp_phylo_with_genus_names_and_BLASTp_bitscore.pdf", width = 30, height = 20)
```

```
Scale for 'colour' is already present. Adding another scale for 'colour',
which will replace the existing scale.

Scale for 'y' is already present. Adding another scale for 'y', which will
replace the existing scale.

Warning message:
“Removed 55 rows containing missing values (geom_image).”
Warning message:
“Removed 55 rows containing missing values (geom_image).”
```

Second, using the E-Value scores:

In [33]:

```
p4 %<+% phylopic_info + geom_nodelab(aes(image=phylopic, colour = phylum), geom="phylopic", alpha=.7) + theme(legend.position = "none")+ 
            scale_colour_manual(values = c("#003b76","#92b74f","#0038d6","#9e5000","#cb54ff",
                                           "#005826","#ff6fe6","#d89f7b","#5a0091","#ff446f",
                                           "#868aff","#3c004c","#aca8c5"),
                                breaks = c('Chordates','Echinoderms','Hemichordates','Platyhelminthes',
                                           'Annelids','Molluscs','Arthropods','Nematodes','Rotifers',
                                           'Cnidaria','Placozoa','Porifera','Ascomycota (Fungi)')) -> p10
options(repr.plot.width = 20, repr.plot.height = 20)
gheatmap(p10, low = "#440154FF", high = "#FDE725FF",font.size = 8,
         odf2_evalue_table, 
         colnames_position = "top", 
         width = 0.25, 
         offset=0.3,
         colnames_angle = 0, 
         legend_title = "-log10(BLASTp\nE-value)", 
         colnames_offset_y = 1, 
         custom_column_labels = c("Cnxn", "Cnxn\nC-ter", "Cnxn\nN-ter")) + ggplot2::ylim(-0.5, NA) + theme(legend.position="bottom")
ggsave("supp_phylo_with_genus_names_and_BLASTp_evalue.pdf", width = 30, height = 20)
```

```
Scale for 'colour' is already present. Adding another scale for 'colour',
which will replace the existing scale.

Scale for 'y' is already present. Adding another scale for 'y', which will
replace the existing scale.

Warning message:
“Removed 55 rows containing missing values (geom_image).”
Warning message:
“Removed 55 rows containing missing values (geom_image).”
```

In [34]:

```
p4 %<+% phylopic_info + geom_nodelab(aes(image=phylopic, colour = phylum), geom="phylopic", alpha=.7) + theme(legend.position = "none")+ 
            scale_colour_manual(values = c("#003b76","#92b74f","#0038d6","#9e5000","#cb54ff",
                                           "#005826","#ff6fe6","#d89f7b","#5a0091","#ff446f",
                                           "#868aff","#3c004c","#aca8c5"),
                                breaks = c('Chordates','Echinoderms','Hemichordates','Platyhelminthes',
                                           'Annelids','Molluscs','Arthropods','Nematodes','Rotifers',
                                           'Cnidaria','Placozoa','Porifera','Ascomycota (Fungi)')) -> p10
options(repr.plot.width = 30, repr.plot.height = 20)
gheatmap(p10, low = "#440154FF", high = "#FDE725FF",font.size = 8,
         odf2_status_table, 
         colnames_position = "top", 
         width = 0.25, 
         offset=0.3,
         colnames_angle = 0, 
         legend_title = "ad-hoc confidence\nassignment", 
         colnames_offset_y = 1, 
         custom_column_labels = c("Cnxn", "Cnxn\nC-ter", "Cnxn\nN-ter")) + ggplot2::ylim(-0.5, NA) + scale_fill_manual(values = c("#71a659","#8975ca","#c5783e","#cb5582"), breaks = c("high conf.", "medium conf.", "low conf.", "unlikely orth."))
ggsave("supp_phylo_with_genus_names_and_BLASTp_confidence_assignment_side_legend.pdf", width = 30, height = 20)
```

```
Scale for 'colour' is already present. Adding another scale for 'colour',
which will replace the existing scale.

Scale for 'y' is already present. Adding another scale for 'y', which will
replace the existing scale.

Scale for 'fill' is already present. Adding another scale for 'fill', which
will replace the existing scale.

Warning message:
“Removed 55 rows containing missing values (geom_image).”
Warning message:
“Removed 55 rows containing missing values (geom_image).”
```
